# A non-linear relation between levels of adult hippocampal neurogenesis and expression of the immature neuron marker doublecortin

**DOI:** 10.1101/2020.05.26.115873

**Authors:** Indira Mendez-David, Denis J David, Claudine Deloménie, Jean-Martin Beaulieu, Alain M. Gardier, René Hen

## Abstract

We investigated the mechanisms underlying the effects of the antidepressant fluoxetine on behavior and adult hippocampal neurogenesis (AHN). After confirming our earlier report that the signaling molecule β2-arrestin is required for the antidepressant-like effects of fluoxetine, we found that the effects of fluoxetine on proliferation of neural progenitors and on survival of adult-born granule cells are absent in the β2-arrestin knockout (β2-Arr KO) mice. To our surprise fluoxetine induced a dramatic upregulation of doublecortin (DCX) in the β2-Arr KO mice, indicating that DCX expression can be increased even though AHN is not. We discovered two other conditions where DCX expression is regulated non linearly compared to levels of AHN: a chronic stress model where DCX is upregulated and an inflammation model where DCX is down regulated. We conclude that assessing DCX expression alone to quantify levels of AHN can be misleading and that caution should be applied when label retention techniques are not available.

**HIGHLIGHTS:**

- β2-arrestin (β-Arr2) is required for the antidepressant-like effects of fluoxetine.
- A dramatic upregulation of doublecortin (DCX) is observed in the β2-Arr KO mice after antidepressant treatment whereas its effects on proliferation of neural progenitors and on survival of adult-born granule cells are absent.
- DCX is more upregulated than the number of young neurons in a mouse model of depression.
- DCX is more down regulated than the number of young neurons in a model of inflammation.
- microRNAs (miRs) may contribute to the regulation of DCX mRNA expression.

## INTRODUCTION

In the adult mammalian brain, there are two regions where new neurons are generated until old age, the subventricular zone (SVZ) which gives rise to neurons that migrate to the olfactory bulb, and the subgranular zone (SGZ) of the hippocampus which gives rise to adult-born granule cells localized in the dentate gyrus (DG). In mice both the SVZ and SGZ neural stem cell niches give rise to new neurons until old age although the numbers decrease significantly with age [for review (Snyder, 2019)]. In primates including humans, the SVZ niche produces young neurons until about one year of age after which neurogenesis becomes undetectable (Sanai et al., 2011); in contrast the SGZ niche keeps producing young neurons considerably later; however there is currently a controversy as to how long the human SGZ keeps producing young neurons. Most reports indicate that adult-born granule cells are generated until old age (Boldrini et al., 2018; Boldrini et al., 2019a; Flor-Garcia et al., 2020; Tartt et al., 2018) but a few reports suggest that the production of young neurons becomes negligible in humans and very low in non-human primates after adolescence (Sorrells et al., 2018). This controversy may be partially due to experimental conditions (such as post-mortem intervals as well as tissue conservation and fixation) (Flor-Garcia et al., 2020) but also to the fact that there are few markers of young neurons that have been studied in the human brain; most studies have used doublecortin (DCX) and as we show in the current study levels of DCX are not always a good estimate of the number of adult-born neurons.

In rodents, adult hippocampal neurogenesis (AHN) has been implicated in a number of cognitive functions as well as in mood and anxiety related behaviors (David et al., 2009; Sahay et al., 2011). In particular, some of the effects of antidepressants such as Selective Serotonin Reuptake Inhibitors (SSRIs) require neurogenesis in the DG (David et al., 2009; Lino de Oliveira et al., 2020; Santarelli et al., 2003). The SSRI fluoxetine has been shown to stimulate several stages of the neurogenesis process: proliferation of the transit amplifying cells, survival of the young adult-born granule cells and differentiation of these young granule cells into mature granule cells (Encinas et al., 2006; Malberg et al., 2000). The mechanisms underlying these effects are only partially understood; activation of several serotonin receptors including the 5-HT_1A_, 5-HT_4_, 5-HT_5A_, and 5-HT_2A_ receptors appears to be important as well as an increase in Brain Derived Neurotrophic Factor and the resulting activation of the Trk-B receptor (David et al., 2009; Kobayashi et al., 2010; Ma et al., 2017; Sagi et al., 2019). There is also some evidence indicating that the signaling molecule β2-arrestin which operates downstream of a number of G-protein coupled receptors (such as 5-HT_4_ and 5-HT_2A_ receptors) is involved in mood disorders and plays a role in the action of antidepressants (Asth et al., 2016; Beaulieu et al., 2008; David et al., 2009). In the current study we investigated the contribution of β2-arrestin to the effects of the antidepressant fluoxetine and discovered a surprising effect of fluoxetine on expression of doublecortin, a commonly used marker for young adult-born granule cells in the DG.

## RESULTS

### 1: The antidepressant/anxiolytic effects of fluoxetine are absent in the β-arrestin-2 heterozygote and Knockout mice

We have been investigating the mechanisms underlying the antidepressant/anxiolytic effects of fluoxetine (18 mg/kg/day) in a variety of animal models. Our previous studies showed that β-arrestin-2 was necessary for several of the behavioral effects of chronic fluoxetine (David et al., 2009). Here we have confirmed these findings by comparing the effects of a 4-week fluoxetine treatment in Wild-Type (WT), β-arrestin2 heterozygote (β-Arr2 Het) and Knockout (β-Arr2 KO) mice in three anxiety/depression tests: the open field (OF), the elevated plus maze (EPM), and the novelty-suppressed feeding test (NSF) **(Figure 1A).** A two-way ANOVA on all anxiety-related parameters revealed that chronic fluoxetine treatment had an effect in WT in the three paradigms **(Supplementary table** 1), resulting in an increase in time spent **(Figure 1B)** and total entries in the center in the OF **(Figure 1C)** without affecting locomotor activity **(Figure 1D),** increase in time spent **(Figure 1E)** and a trend for entries **(Figure 1F)** in open arms in the EPM, a trend for an decrease in latency to feed in the NSF **(Figure 1G)** without affecting food consumption **(Figure 1H).** All these fluoxetine-induced anxiolytic/antidepressant effects are absent in β-Arr2 KO mice **(Figure 1B–1E).** Interestingly, these anxiolytic/antidepressant effects of fluoxetine are also absent in the β-Arr2 Het mice, which indicate that both alleles of β-Arrestin2 are necessary for the effects of fluoxetine. In keeping with our previous findings (David et al., 2009) the β-Arr2 KO display more anxiety-related behaviors than their WT littermates as shown by a decrease in time spent in the center and entries in the center of the OF, a trend for a decrease in time spent in the open arms of the EPM and an increase in latency to feed in the NSF **(Figure 1B–1E).** In addition, we show here that the β-Arr2 Het mice display an intermediate phenotype between the WT and the KOs in most anxiety-related values **(Figure 1B–1E).**

**Figure 1:**
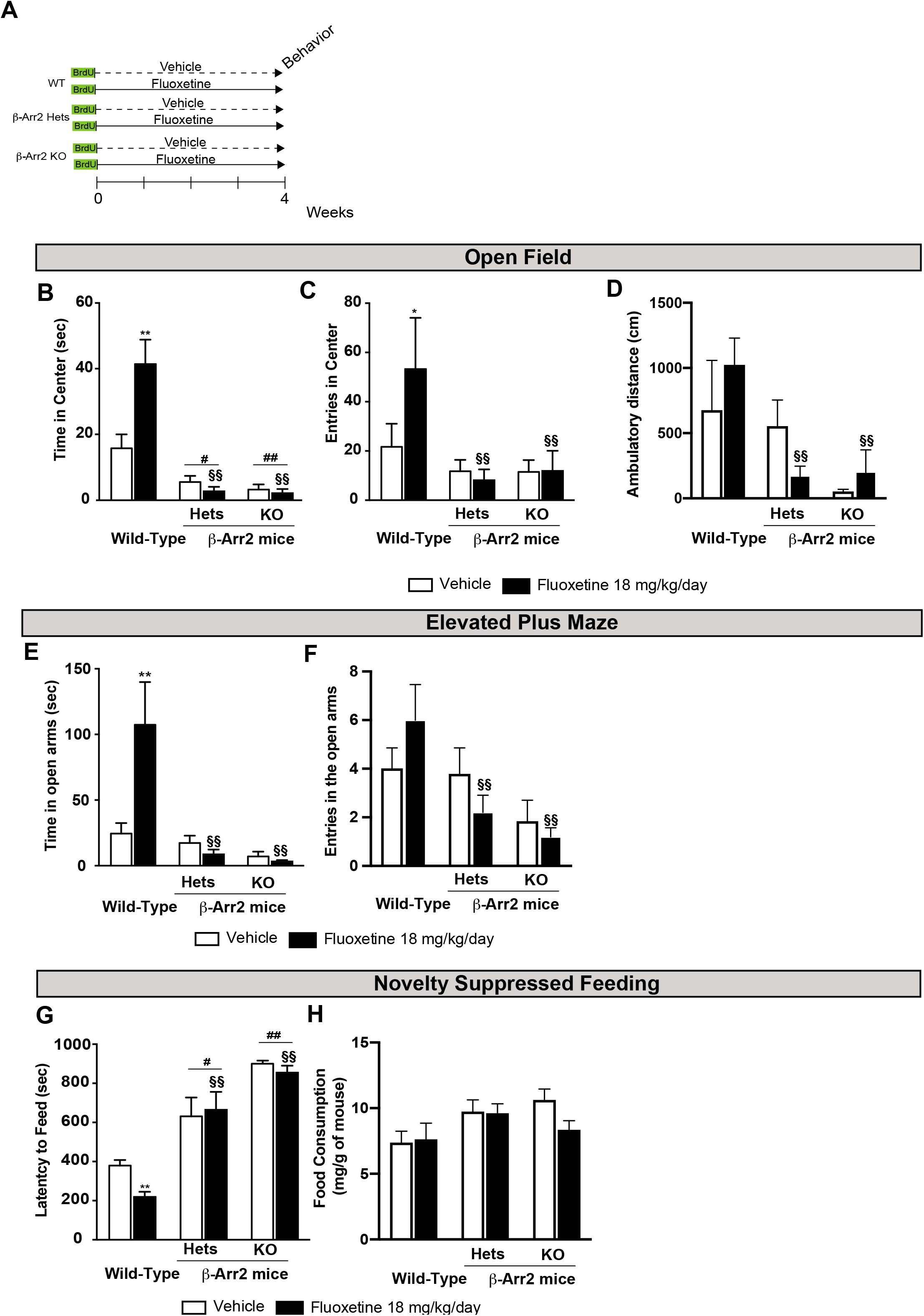
β-arrestin-2 expression is involved in chronic fluoxetine induced anxiolytic/antidepressant-like effects. (A) Experimental protocol timeline to assess behavioral and neurogenic consequences of a 4-week fluoxetine treatment (18 mg/kg/day in the drinking water) in β-arrestin-2 Heterozygous (β-Arr2 Hets), knockout mice (β-Arr2 KO) mice compared with their Wild-Types littermates (WT). (B-H) To evaluate survival of newborn cells, 5-Bromo-2-Deoxyuridine (BrdU) was administered twice a day at 150 mg/kg intraperitoneally, during 3 days, 5 weeks before sacrifice. Anxiety was expressed as the time spent in the center (B), the total entries in the center (C), the total ambulatory distance (D) during 30 min in the open field and as time spent (E) or entries (F) in open arms of the elevated plus maze. In the Novelty Suppressed Feeding, latency time to feed was measured (G) and food consumption in home cage (H). Values plot are mean ± SEM (n = 4–8 animals/group). Data were analyzed with a two-way ANOVA (see supplementary table 1). Significant main effects and/or interactions were followed by Fisher’s PLSD post-hoc analysis (see supplementary table 1). *p< 0.05, **p< 0.01 for comparisons between vehicle-treated WT and fluoxetine-treated WT mice; #*p*< 0.05, ##p< 0.01 for comparisons between fluoxetine-related β-Arr2 Hets or β-Arr2 KO and vehicle-treated WT mice; §§p< 0.01 comparisons between fluoxetine-treated β-Arr2 Hets or β-Arr2 KO and fluoxetine-treated WT mice.

### 2: The effects of fluoxetine on neurogenesis are absent in the β-arrestin-2 Heterozygous and Knockout mice but paradoxically the expression of two markers of immature neurons, doublecortin and calretinin, is dramatically upregulated by fluoxetine in KO mice

Since AHN has been implicated in some of the effects of antidepressants (David et al., 2009; Lino de Oliveira et al., 2020; Santarelli et al., 2003) we also examined the effects of chronic fluoxetine treatment on various stages of the neurogenesis process in the β-Arr2 Het and β-Arr2 KO mice. Proliferation in the SGZ was assessed by the level of KI67 a marker of dividing cells; survival of the young neurons was assessed by retention of the BrdU nucleotide analog injected 5 weeks prior to sacrifice; and numbers of immature neurons were assessed by immunocytochemistry against two markers of immature neurons: DCX and calretinin (CR) which are expressed during the first month of the maturation of young adult-born neurons (Kempermann et al., 2015).

A two-way ANOVA revealed that the stimulatory effect of fluoxetine on proliferation and survival observed in WT mice **(Figure 2A–2B)** was absent in β-Arr2 Het and KO mice **(Supplementary table 2).** These results mirror the behavioral effects of fluoxetine and suggest that the anxiolytic/antidepressant-like effects of fluoxetine may be mediated at least in part by an increase in neurogenesis.

**Figure 2:**
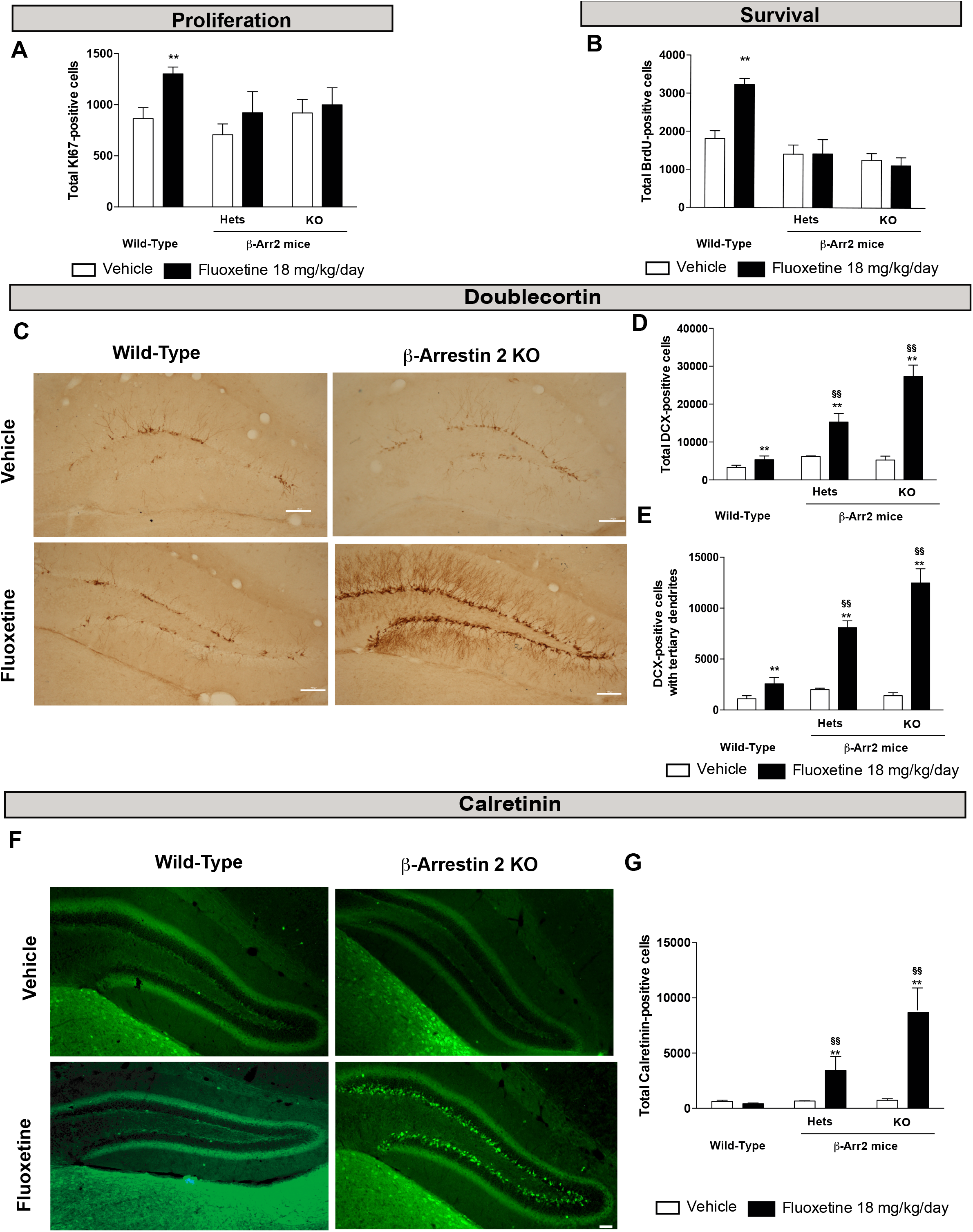
β-arrestin-2 expression is required for chronic fluoxetine treatment increased survival of newborn cells but not maturation of young neurons. (A-G) The effects of 28 days of treatment with fluoxetine (18 mg/kg/day) on cell proliferation (A), cell survival (B) and maturation of young neurons (C-G) were compare to those of vehicle in β-arrestin-2 Heterozygous (β-Arr2 Hets), knockout mice (β-Arr2 KO) mice compared with their Wild-Types littermates (WT). Proliferation (A) and survival (B) are measured as Ki67+ and BrdU+ cells respectively. Maturation was characterized by the total number of DCX+ cells (D), the number of DCX+ cells with tertiary dendrites (E) and calretinin+ cells (G). Images of doublecortin or calretinin staining following vehicle or fluoxetine treatment in β-arrestin-2 KO mice and their WT littermates (10x magnification, scale bar= 100 μm) (C, F). Values plot are mean ± SEM (n = 3-7 animals/group). Data were analyzed with a two-way ANOVA (see supplementary table 2). Significant main effects and/or interactions were followed by Fisher’s PLSD post-hoc analysis (see supplementary table 2). *p< 0.05; **p< 0.01 for comparisons between vehicle-treated group and fluoxetine treated group for each genotype; §§p< 0.01 comparisons between fluoxetine-treated β-Arr2 Hets or β-Arr2 KO and fluoxetine-treated WT mice.

Surprisingly, the effects of fluoxetine on the expression of two markers of immature neurons, DCX and CR were dramatically different. A two-way ANOVA revealed a massive upregulation of DCX and CR in the β-Arr2 KO mice **(Figure 2C–2G)** even though these mice do not display any increase in neurogenesis as measured by KI67 and BrdU incorporation **(Figure 2A–2B).** This upregulation of DCX and CR appears to be dependent on the expression level of β-arrestin-2 because it is intermediate in the β-Arr2 Het mice **(Figure 2C–2G).**

This is to our knowledge the first example of a complete dissociation between levels of neurogenesis and levels of DCX; namely in the β-Arr2 Het and KO mice where neurogenesis is not increased by fluoxetine neither at the level of proliferation in the SGZ nor at the level of survival of the young neurons, DCX and CR levels are massively upregulated. This suggests that the levels of these proteins can be increased without a change in neuronal number and that this upregulation only happens when β-arrestin-2 levels are decreased.

Since two types of β-arrestins are expressed in the hippocampus (David et al., 2009), β-arrestin-1 and β-arrestin-2, we conducted a similar study in the β-arrestin-1 (β-Arr1) KO mice **(Supplemental Figure 1, Supplementary table 3).** Like in the β-Arr2 KO proliferation and survival were not increased by chronic fluoxetine in the β-Arr1 KO **(Supplemental Figure 1A-1C).** In contrast DCX was increased to the same extent in β-Arr1 KO as in the WT **(Supplemental Figure 1D-1F).** Therefore, we also see a disconnect between expression of DCX and levels of neurogenesis in the β-Arr1 KO but to a lesser extent than in the β-Arr2 KO.

We also analyzed the expression of DCX in the piriform cortex of β-Arr2 Het, KO and their littermates **(Supplemental Figure 2, Supplementary table 4)** because this is one of the few brain areas where DCX expression has been observed (Klempin et al., 2011). We found an increase in total DCX and DCX with tertiary dendrites expression in β-Arr2 KO mice **(Supplemental Figure 2A-C).** We also found an elevated basal level of DCX in the β-Arr2 KO compared to the WT. However, no BrdU positive cells were found (data not shown) confirming previous report that DCX-expressing cells in the piriform cortex were strictly postmitotic (Klempin et al., 2011). Therefore, this is an instance where DCX can be detected and regulated in a non-neurogenic niche, the piriform cortex (Kremer et al., 2013). Interestingly, in both the hippocampus and piriform cortex β-arrestin-2 signaling appears to suppress DCX expression.

### 3: In the corticosterone model of anxiety/depression, fluoxetine induces a larger increase in the expression of DCX and Calretinin than in levels of neurogenesis

We next decided to assess whether a dissociation between neurogenesis and DCX levels can also be found in other situations than in the β-Arr1 or β-Arr2 KO mice. We chose to look at a model of anxiety/depression, the chronic corticosterone (CORT) model **(Figure 3),** because a few previous studies had shown surprisingly large effects of fluoxetine on DCX levels in this model (David et al., 2009; Mendez-David et al., 2014; Robinson et al., 2016). This model consists in administering CORT for 4 weeks in the drinking water, which produces an increase in anxiety, and depression-related measures, that can be reversed by a chronic antidepressant treatment (David et al., 2009; Mendez-David et al., 2014; Mendez-David et al., 2017) **(Figure 3A).** In control animals that did not receive CORT, fluoxetine elicited a similar modest increase in survival (+11%), and levels of DCX (+24%) or DCX with tertiary dendrites (+26%) **(Figure 3B-E, Supplementary table** 5). In contrast, in CORT-treated animals, the increase in DCX level after chronic fluoxetine was significantly larger than the increase in survival **(Figure 3C-D).** This effect was more pronounced when we counted the number of DCX positive cells with tertiary dendrites (+199%), which are the most differentiated cells (**Figure 3E**). When we analyzed another marker of immature neurons, calretinin, the effect of fluoxetine in the CORT group was even more dramatic: 12-fold [567 ±62 vs 6834 ±1234 in CORT/vehicle and CORT/fluoxetine respectively **(Figure 3F-G)** while the survival was only increased by 22% (1272 ±68 vs 1558 ±115 in CORT/vehicle and CORT/fluoxetine respectively).

**Figure 3:**
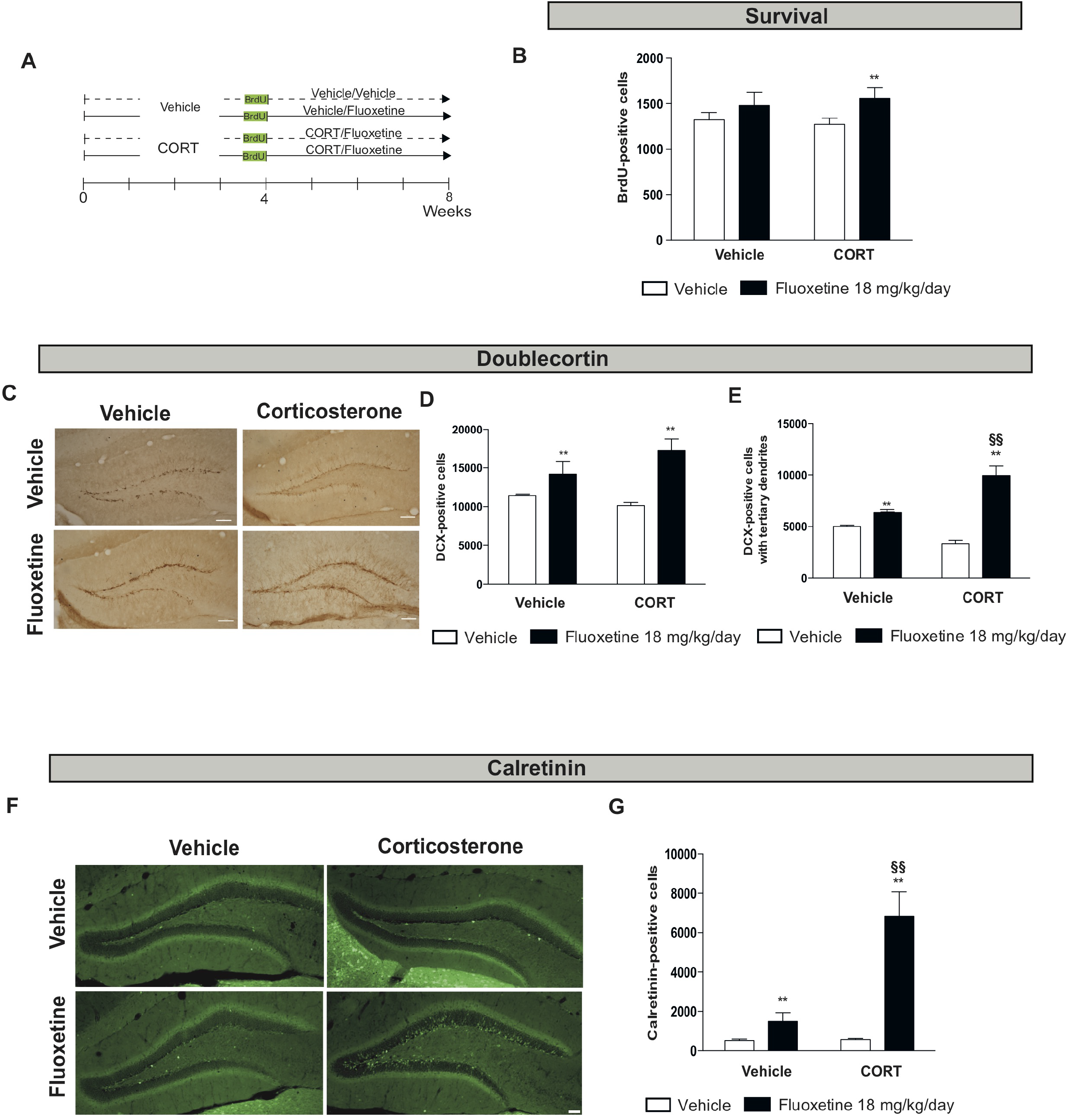
Chronic Fluoxetine treatment-increased doublecortin and calretinin expression in adult dentate gyrus independently from adult hippocampal neurogenesis. (A) Experimental protocol timeline to assess survival and dendritic maturation of young neurons in the dentate gyrus of the hippocampus of 8 weeks of corticosterone (35 μg/ml in the drinking water) ± fluoxetine (18 mg/kg/day in the drinking water) during the last 4 weeks. To evaluate survival of newborn cells, 5-Bromo-2-Deoxyuridine (BrdU) was administered twice a day at 150 mg/kg, intraperitoneally, for 3 days 4 weeks before sacrifice. (B-G) The effects of 28 days of treatment with fluoxetine (18 mg/kg/day) on cell survival (B) and maturation of young neurons (C-G) were compare to those of vehicle in corticosterone-treated mice or not. Survival (B) was measured BrdU+ cells. Maturation was characterized by the total number of DCX+ cells (D), the number of DCX+ cells with tertiary dendrites (E) and calretinin+ cells (G). Images of doublecortin or calretinin staining following vehicle or fluoxetine treatment in corticosterone-treated animals or not. 10x magnification (scale bar= 100 μm) (C, F). Values plot are mean ± SEM (n = 3-5 animals/group). Data were analyzed with a two-way ANOVA (see supplementary table 5). Significant main effects and/or interactions were followed by Fisher’s PLSD post-hoc analysis, **p< 0.01 for comparisons between fluoxetine-treated groups and vehicle-treated groups in control (Vehicle) or CORT mice; §§p< 0.01 comparisons between fluoxetine-treated CORT groups and fluoxetine treated control groups.

The transition from immature to mature neuronal markers is characterized by a higher order dendritic branching (tertiary and higher) as well as a partial migration from the SGZ into the granule cell layer (GCL) in mice (Klempin et al., 2011; Kohler et al., 2011) and in non-human primates (Ngwenya et al., 2015). Therefore, we also examined this migration of DCX positive cells in the CORT model after chronic fluoxetine by counting the proportion of DCX+ cells within the GCL. Like in the case of tertiary dendrites we found a large increase in the CORT + fluoxetine group **(Supplemental Figure 3, Supplementary table 6).**

In aggregate, these results indicate again a dissociation between levels of neurogenesis and expression of DCX and CR, in this case after a chronic treatment with corticosterone.

### 4: The ablation of neurogenesis prevents the effect of fluoxetine on DCX and calretinin expression

An earlier report (Kobayashi et al., 2010) and a recent review (Ohira et al., 2019) had suggested that fluoxetine induced, in addition to neurogenesis, a dematuration of mature granule cells that would lead to the expression of immature markers such as DCX and Calretinin. We wondered therefore whether such a dematuration might contribute to the increase in DCX and CR elicited by fluoxetine in the CORT model. To test for that hypothesis, we ablated neurogenesis using two different approaches, one with X-irradiation and another one with a pharmacogenetic strategy prior to assessing the impact of fluoxetine on DCX and CR expression in the CORT paradigm **(Figure 4).** This pharmacogenetic strategy utilizes the glial fibrillary acidic protein (GFAP) promoter-driving the Herpes Simplex Virus Thymidine Kinase (TK) gene. As a result, the dividing GFAP positive progenitor cells die following treatment with the antiviral drug valganciclovir (vGCV) (Mendez-David et al., 2017; Saxe et al., 2006) **(Figure 4C).** We found no detectable expression of DCX after X-irradiation or in the GFAP-TK+ mice treated with chronic fluoxetine **(Figure 4B-D).** These results indicate that mature granule cells which are spared in this ablation model, do not contribute significantly to the observed increase in DCX. As previously shown (Klempin et al., 2011), we also showed that most DCX+ cells co-expressed calretinin another marker of immature neurons and that X-irradiation suppressed the expression of both DCX and calretinin **(Supplemental Figure 4, Supplementary table 7).** We conclude therefore that the increase in DCX and calretinin levels observed after fluoxetine in the CORT-treated group comes mostly from immature adult-born neurons rather than from a dematuration of mature granule cells.

**Figure 4:**
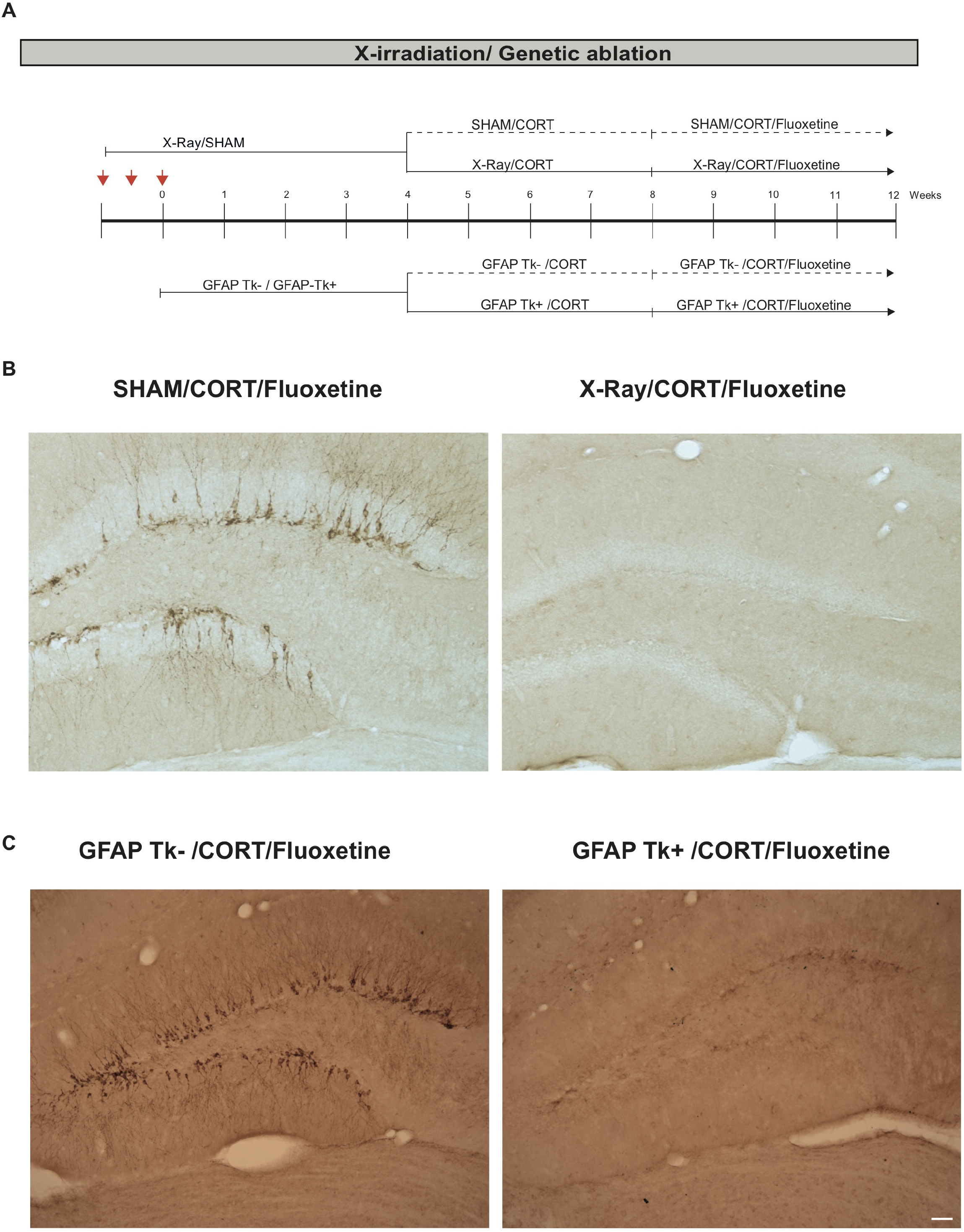
Doublecortin expression is sensitive to adult hippocampal neurogenesis ablation. (A) Experimental protocol timeline to assess consequences of arresting adult hippocampal neurogenesis using focal X-irradiation or pharmacogenetic in chronic fluoxetine –treated mice (18 mg/kg/day for 28 days in the drinking water) in a stress-related model of Anxiety/Depression after 4 weeks of corticosterone (35 μg/ml in the drinking water). X-irradiation occurred a month before the start of corticosterone treatment to ensure that markers of inflammation are indistinguishable from sham animals. Genetic ablation was performed using GFAP TK+ mice receiving vGCV a month before the start of corticosterone. (B) Images of doublecortin staining following chronic fluoxetine treatment in X-irradiated or sham corticosterone-treated animals (10x magnification). (C) Images of doublecortin staining following chronic fluoxetine treatment in GFAP-TK+ or GFAP-TK-mice under corticosterone treatment (10x magnification, scale bar= 100 μm).

### 5: Inflammation induces a rapid decrease in DCX expression

We decided also to assess the expression of DCX after inflammation because our preliminary result had indicated that DCX levels decreased dramatically immediately after X-irradiation event though the number of young neurons was not much affected (Burghardt et al., 2012; Denny et al., 2012). To induce inflammation in a controlled way we performed intrahippocampal administration of Lipopolysaccharide (LPS). LPS is an endotoxin of gram-negative bacteria, used extensively for inducing an immune response. Unilateral injection of LPS in the dorsal DG were performed at and only the right side of the brain and mice were sacrificed at three different time points after LPS injection: 24h, 3 and 7 days; therefore, the vehicle-injected side and the ventral pole of the DG can be used as controls **(Figure 5A).** The number of young neurons as measured by BrdU incorporation is not significantly decreased after 24 hrs of LPS injection **(Figure 5B, Supplementary table 8).** In contrast, in keeping with our previous X-ray data there is a marked decrease in DCX expression 24 hours after LPS injection and only on the injected side and not on the ventral pole **(Figure 5C-D).** These results suggest that like X-irradiation, LPS-induced inflammation results in a decrease in DCX expression in the absence of a significant change in the number of young adult-born neurons. 3 and 7 days after LPS injection the decrease of DCX expression remains restricted to the injected side and a decrease in the number of young neurons (BrdU labelled) becomes visible although it is less pronounced than the decrease in DCX expression **(Figure 5B-D).** In keeping with these results the fraction of BrdU positive cells that are also DCX positive is larger on the control than on the LPS-injected side at all three time points **(Figure 5E).**

**Figure 5:**
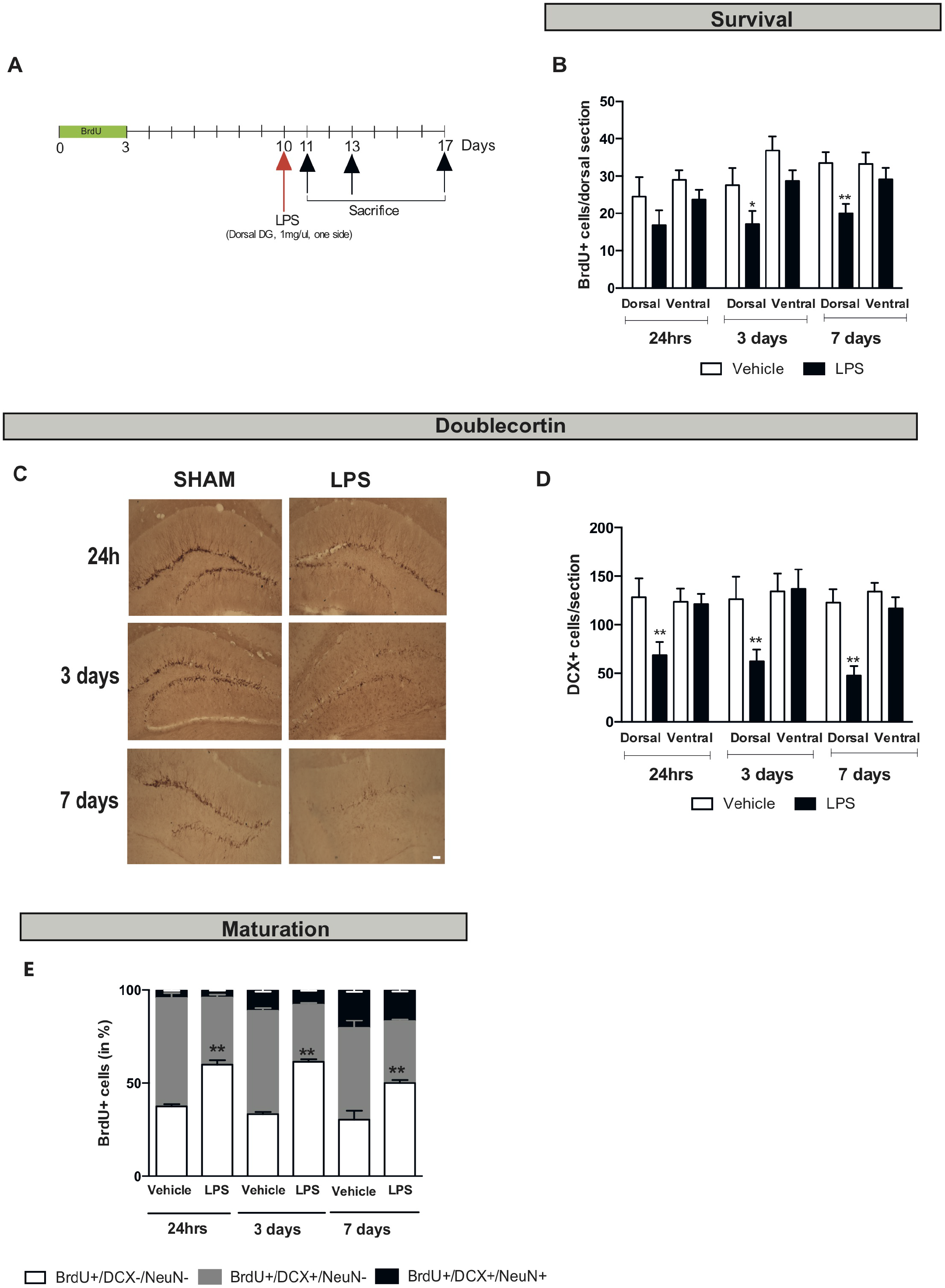
Injecting a pro-inflammatory lipopolysaccharide in the mouse dorsal dentate gyrus, induced a large reduction in doublecortin expression. (A) Experimental protocol timeline to assess consequences of a unilateral lipopolysaccharide (LPS) infusion (1mg/μl) in the dorsal dentate gyrus on survival and dendritic maturation of young neurons. To evaluate survival of newborn cells, 5-Bromo-2-Deoxyuridine (BrdU) was administered twice a day at 150 mg/kg intraperitoneally for 3 days, 7,10 or 14 days before sacrifice. (B-E) The effects of LPS-induced inflammation on survival of newborn cells (B), DCX+ cells (C-D) or maturation of newborn neurons were quantified by the number of Brdu+ cells in the dorsal dentate gyrus at the injected side (right side, R) and compare to the control-lateral side (left side, R) 24h, 3 days and 7 days post-injection along the dorso/lateral axis (E). Images of doublecortin staining (C) following LPS infusion at 24h, 3 days and 7 days compared to the vehicle-injected side (10x magnification, scale bar= 100 μm). The effect of LPS on DCX+ cells was measured at the injected (R) along the dorso-lateral axis, 24h, 3 days and 7 days post injection when compared and the controllateral vehicle-injected side (L) (D). The fate of surviving BrdU+ was investigated by triple staining using co-localization of BrdU+ cells with markers of neuronal maturation (DCX) and mature neurons (NeuN) 24h, 3 days and 7 days post LPS injection (F). Values plot are mean mean ± SEM (n = 3–7 animals/group). Data were analyzed with a one-way ANOVA with repeated measure (see supplementary table 8). Significant main effects and/or interactions were followed by Fisher’s PLSD post-hoc analysis (see supplementary table 4) *p< 0.05; **p< 0.01 for comparisons between the LPS-injected and the vehicle-injected side along the dorsal/ventral axis

In summary we have shown that the expression of DCX can be upregulated in young adult-born neurons (fluoxetine in β-Arr-2 or β-Arr-1 KO mice and in CORT model) or down regulated (inflammation) independently of the number of these young neurons.

### 6: Fluoxetine modulates the expression of MIRs that have been implicated in DCX expression

In order to start assessing the mechanism that may regulate expression of DCX and CR in response to fluoxetine in the CORT model, we investigated the level of expression of several microRNAs (miR) that have been implicated in the regulation of DCX mRNA expression **(Figure 6A).** First, we showed that levels of DCX mRNA are upregulated by fluoxetine in the CORT model by RT-PCR **(Figure 6B).** Next, we investigated the levels of miR-128, miR-18a5, miR-22a3, and miR-22a5 by RT-PCR **(Figure 6C-F, Supplementary table 9).** We chose miR128 because it had been shown to downregulate the expression of DCX by directly binding on its 30□TR region (Evangelisti et al., 2009), and miR-18a, miR-22a3, and miR-22a5 because they belong to a cluster shown to regulate neurogenesis as well as anxiety and depression related behaviors (Dwivedi, 2013; Jin et al., 2016). Chronic fluoxetine treatment resulted in a trend for a decrease in miR-128 expression **(Figure 6C).** In contrast, chronic fluoxetine significantly increased the levels of miR-18a5, miR-22a3 **(Figure 6D-E).** In regard to miR-22a5, a trend for an increase was also observed **(Figure 6D-E).** These results are consistent with the previously described impact of these miRs on DCX expression and neurogenesis. It is therefore possible that these miRs contribute to the regulation of DCX mRNA expression in our fluoxetine group as well as possibly in our other experimental manipulations such as the β-Arr-2 KO and the LPS-induced inflammation.

**Figure 6:**
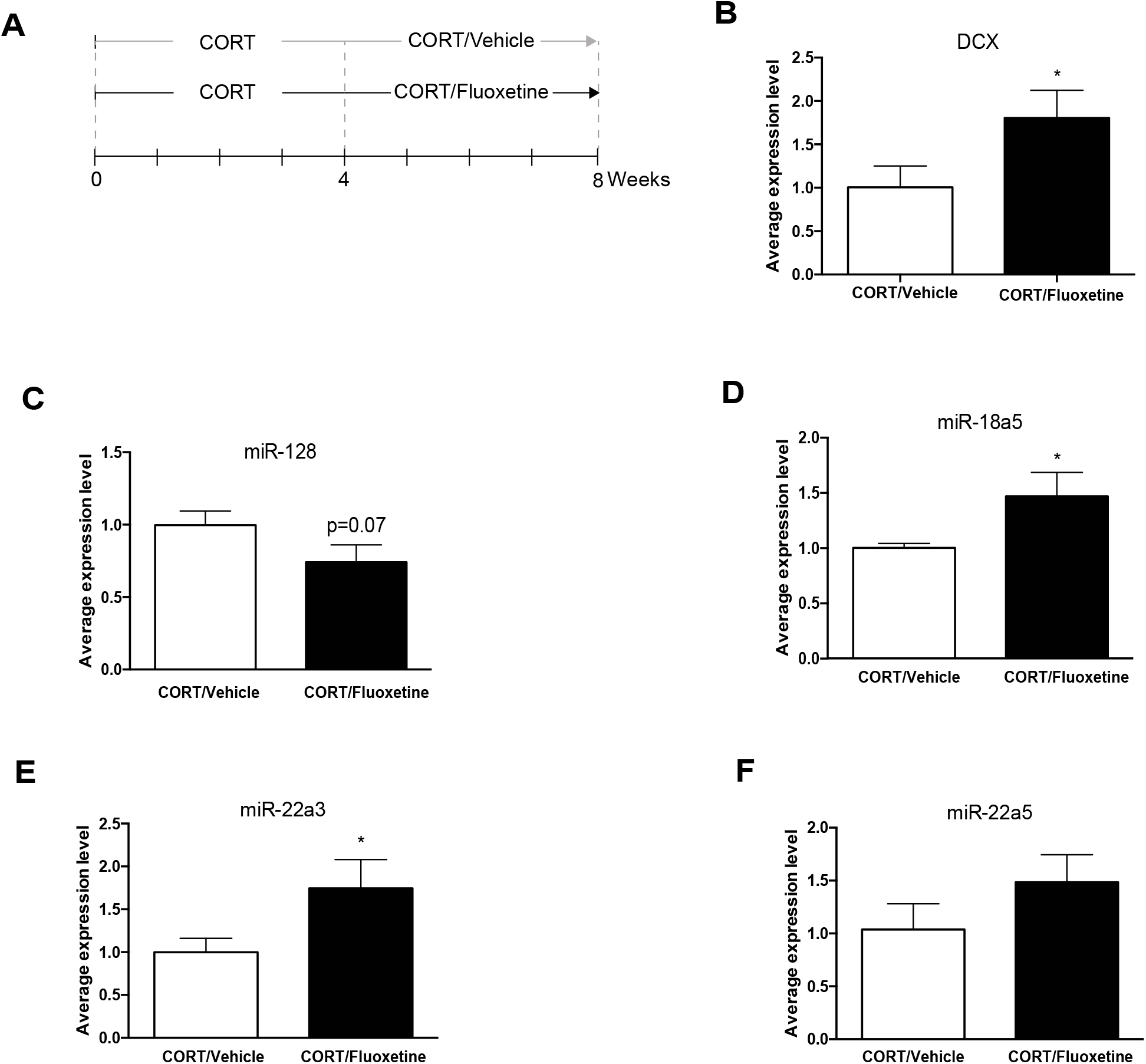
Chronic fluoxetine –increased hippocampal doublecortin expression in a stress-related model of Anxiety/Depression modulate the expression of microRNA. (A) Experimental protocol timeline to assess the effect of chronic fluoxetine treatment (18 mg/kg/day in the drinking water) in a stress-related model of Anxiety/Depression after 4 weeks of corticosterone (CORT, 35 μg/ml in the drinking water) on hippocampal DCX and microRNA (miR) expression. (B-F) Values plot are mean ± SEM (n= 4-5 animals /group). Data were analyzed with an unpaired t-test (see supplementary table 9). *p< 0.05 unpaired t tests between CORT/fluoxetine and CORT/vehicle-treated groups.

## DISCUSSION

We will briefly discuss the mechanisms that may be responsible for the disconnect between levels of the DCX protein and levels of neurogenesis. We have observed in several models, before enlarging the discussion to the current controversy about levels of neurogenesis in the adult human brain and how our current results may inform that controversy.

### 1: Disconnect between DCX and neurogenesis

The effects of chronic fluoxetine on both anxiety/depression related behaviors and on hippocampal neurogenesis have been shown to be mediated by several 5-HT receptors including the 5-HT_1A_ receptor (Samuels et al., 2016; Santarelli et al., 2003), the 5-HT_4_ receptor (Kobayashi et al., 2010; Mendez-David et al., 2014) and possibly the 5-HT_2B_ (Diaz et al., 2012), 5-HT_1B_ (Svenningsson et al., 2006), 5-HT_2C_ (Opal et al., 2014) and 5-HT_5A_ receptors (Sagi et al., 2019). In particular in the case of the 5-HT_1A_ receptor we have previously shown that 5-HT_1A_ receptors located in mature granule cells of the DG are critical for the antidepressant/anxiolytic effects of chronic fluoxetine and contribute to the effects of fluoxetine on neurogenesis. There is also evidence that the 5-HT_4_ receptor is involved in both the behavioral and neurogenic effects of fluoxetine, but it is not known whether it is the DG 5-HT_4_ receptors that contribute to these effects. Both 5-HT_1A_ and 5-HT_4_ receptors have been shown to engage the β-arrestin signaling pathway (Bohn and Schmid, 2010; Pytka et al., 2018). It is therefore conceivable that the lack of a behavioral response to fluoxetine in the β-Arr2 KO mice is due to a lack of β-arrestin 2 in the DG and a subsequent inability of the 5-HT_1A_ and 5-HT_4_ receptors to engage β-Arr2 signaling pathways such as the MAPK/ERK signaling. Such impairment may also be related to the absence of the effect of fluoxetine on proliferation and survival in the β-arrestin-2 KO mice. What is the most surprising is that the levels of DCX and calretinin, which are both markers of immature neurons, are dramatically upregulated by fluoxetine in the β-arrestin-2 KO mice. Such an observation may be related to the fact that fluoxetine has been shown to accelerate the maturation of young neurons via a faster transition from the DCX positive stage to the NeuN positive stage (Sorrells et al., 2018; Wang et al., 2008). It is possible that in the absence of β-arrestin 2 this transition does not happen or happens more slowly resulting in an accumulation of DCX in the young neurons.

In order to investigate mechanisms that may contribute to the regulation of expression of DCX we analyzed the expression of a number of miRs that had been implicated in stress or the modulation of neurogenesis [for review (Dwivedi, 2014)]. We find that a chronic fluoxetine treatment modulates the expression of these miRs in a direction, which is consistent with the upregulation of DCX mRNA expression by fluoxetine. Such results suggest that these miRs may be responsible at least in part for the regulation of DCX that we observe in our models.

We go on to demonstrate that DCX can also be regulated non linearly compared to levels of neurogenesis in two other models. The chronic CORT model where chronic fluoxetine results in a larger increase in DCX expression than the number of proliferating precursors as measured by KI67 and the number of surviving neurons as measured by BrdU incorporation. Such an observation is in good agreement with an independent report that reached a similar conclusion (Robinson et al., 2016). The other model where we observe a difference between DCX levels and levels of neurogenesis is a model of inflammation where DCX levels decrease more and faster than neurogenesis. Such observations confirm our previous findings that levels of DCX decrease dramatically already 24h after X-irradiation, which induces an acute inflammatory response (Denny et al., 2012) even though most young neurons are not affected by X-irradiation.

Another independent line of evidence showing a complete disconnect between levels of doublecortin and levels of neurogenesis comes from the DCX KO mice that display normal levels of neurogenesis in the absence of any DCX expression (Dhaliwal et al., 2015). Finally, there is evidence that maturation of young neurons is partially independent of the proliferation and survival of these cells (Baptista and Andrade, 2018; Kempermann et al., 2015; Plumpe et al., 2006). For example, in rats levels of DCX are lower than in mice although levels of neurogenesis are higher and this discrepancy may be due to a faster maturation of the young neurons in rats than in mice (Snyder et al., 2016).

However, it is important to emphasize that DCX is still exclusively expressed by immature adult-born neurons since expression of DCX is abolished by two manipulations that suppress adult hippocampal neurogenesis: X-ray and GFAP-TK. These results argue against the possibility that DCX in the dentate gyrus could also come from non-neurogenesis related processes as has been argued by some authors (Kobayashi et al., 2010).

### 2. Human neurogenesis

Recent reports have questioned whether AHN occurs in the adult human hippocampus and to what extent (Boldrini et al., 2018; Boldrini et al., 2019b; Moreno-Jimenez et al., 2019; Sorrells et al., 2018; Tartt et al., 2018; Toda et al., 2019). However, most of these studies rely primarily on DCX expression to quantify levels of neurogenesis, which as we just discovered can be problematic. In species where it is possible it is clearly better to use a combination of markers that encompass the different stages of the neurogenesis process: proliferation, maturation and survival. The problem is the most acute in humans where it is difficult if not impossible to assess survival of young neurons because lineage studies and label retention studies with BrdU or retroviruses are mostly impossible. Therefore, most human studies have relied on a few markers of immature neurons such as DCX to quantify neurogenesis. For example, in the DG of adult healthy or Alzheimer patients, (Moreno-Jimenez et al., 2019) observed a relative abundance of DCX+ immature neurons. In contrast Sorrells et al found rare DCX+ cells by 7 and 13 years of age which led these authors to suggest that hippocampal neurogenesis ends in childhood (Sorrells et al., 2018). This problem of quantification is complicated by the fact that these groups used different fixing and staining conditions. In addition, post mortem intervals were highly variable which has been shown to impact DCX expression (Verwer et al., 2007). Finally, levels of stress, inflammation or other health conditions that may impact neurogenesis are not always well known prior to the death of the individual. As we have shown in the current study these factors may have large impacts on DCX expression and may explain at least in part the current controversy about levels of neurogenesis in the adult human brain (Boldrini et al., 2018; Kempermann et al., 2015; Sorrells et al., 2018; Spalding et al., 2013).

In summary, we have shown that DCX expression can be modulated non linearly compared to levels of neurogenesis. Although the functional significance of this regulation remains unknown, we hope that our study will serve as a cautionary note for those interested in quantifying neurogenesis particularly in humans and will emphasize the need for multiple markers to be able to assess the various stages of the neurogenesis process.

## STARE METHODS

Detailed methods are provided in the online version of this paper and include the following:

- **KEY RESOURCES TABLE**
- **CONTACT FOR REAGENT AND RESOURCE SHARING**
- **EXPERIMENTAL MODEL AND SUBJECT DETAILS**
- **METHOD DETAILS**

- **Animals**

*- Study in β-arrestin 2 KO mice*
*- Study in CORT model*
*- Study in irradiated mice*
*- Study in GFAP-TK+ mice*
*- Lipopolysaccharide study*
- **Drugs and treatments**

*- Study in β-arrestin 2 KO mice*
*- Study in CORT model*
*- Arresting adult hippocampal neurogenesis* *X-irradiation *Genetic ablation of adult hippocampal neurogenesis study
*- Lipopolysccharide study*
- **Behavioral study in β-arrestin 2 KO mice**

*- Open Field (OF)*
*- Elevated Plus Maze-Novelty Suppressed Feeding (NSF)*
- **Immunochemistry**

*- Proliferation study*
*- Survival study*
*- Doublecortin (DCX) labelling*
*- Calretinin (CF) labelling*
*- Immunohistochemistry and confocal imaging for maturation study*
- **Hippocampal doublecortin and MicroRNA expression**

*- Hippocampal microdissection and preparation of total RNA*
*- RT-qPCR analysis*
- **STATISTICAL ANALYSIS**

## ACKNOWLEDGEMENTS

We also thank L. Tritschler for his technical contribution in LPS infusion (CESP, Univ Paris-Saclay), I. Pavlova and the C. Denny’s lab for confocal acquisition (Columbia Univ.), V. Domergue and the staff of the animal care facility [UMS-IPSIT (Plateforme Anime au Service de l’Innovation Thérapeutique), Université Paris-Saclay, Châtenay-Malabry, France] for their technical support. We thank M. Pitel [UMS-IPSIT (Ingénierie et Plateformes au Service de l’Innovation Thérapeutique), Université Paris-Saclay, Châtenay-Malabry, France] for her technical support on qPCR. This work was supported by a National Alliance for Research on Schizophrenia and Depression (NARSAD) 2017 Young Investigator award from Brain & Behavior Research Foundation and Deniker Foundation (I.M.D).

## AUTHOR CONTRIBUTION

R.H, A.M.G, I.M.D, D.J.D conceived, planned, oversaw the study. J.M.B. provided key reagents (β-arrestin 1 +/− line). I.M.D, D.J.D, C. D, performed the experiments. I.M.D, D.J.D, C.D. analyzed the data. R.H, I.M.D, D.J.D wrote the paper. A.M.G, C.D, J.M.B. assisted in revising the manuscript. I.M.D and DJD obtained funding and resource used in the experiments.

## DECLARATION OF INTERESTS

DJD serves as a consultant for Lundbeck Inc. DJD receives compensation from Lundbeck and Janssen. RH receives compensation as a consultant for Roche, Lundbeck and Servier in relation to the generation of novel antidepressants.

## STAR□ METHODS

### KEY RESOURCESTABLE

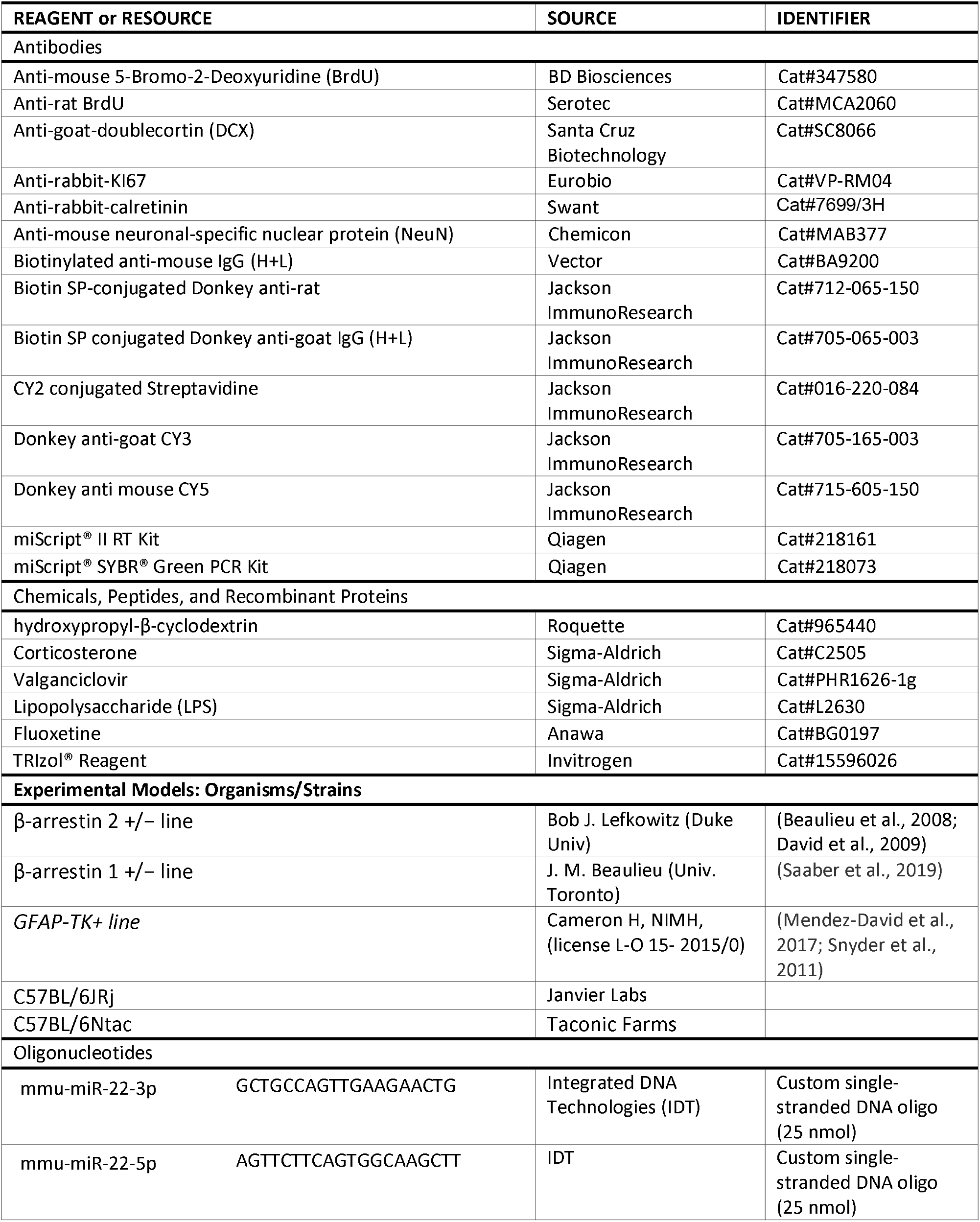
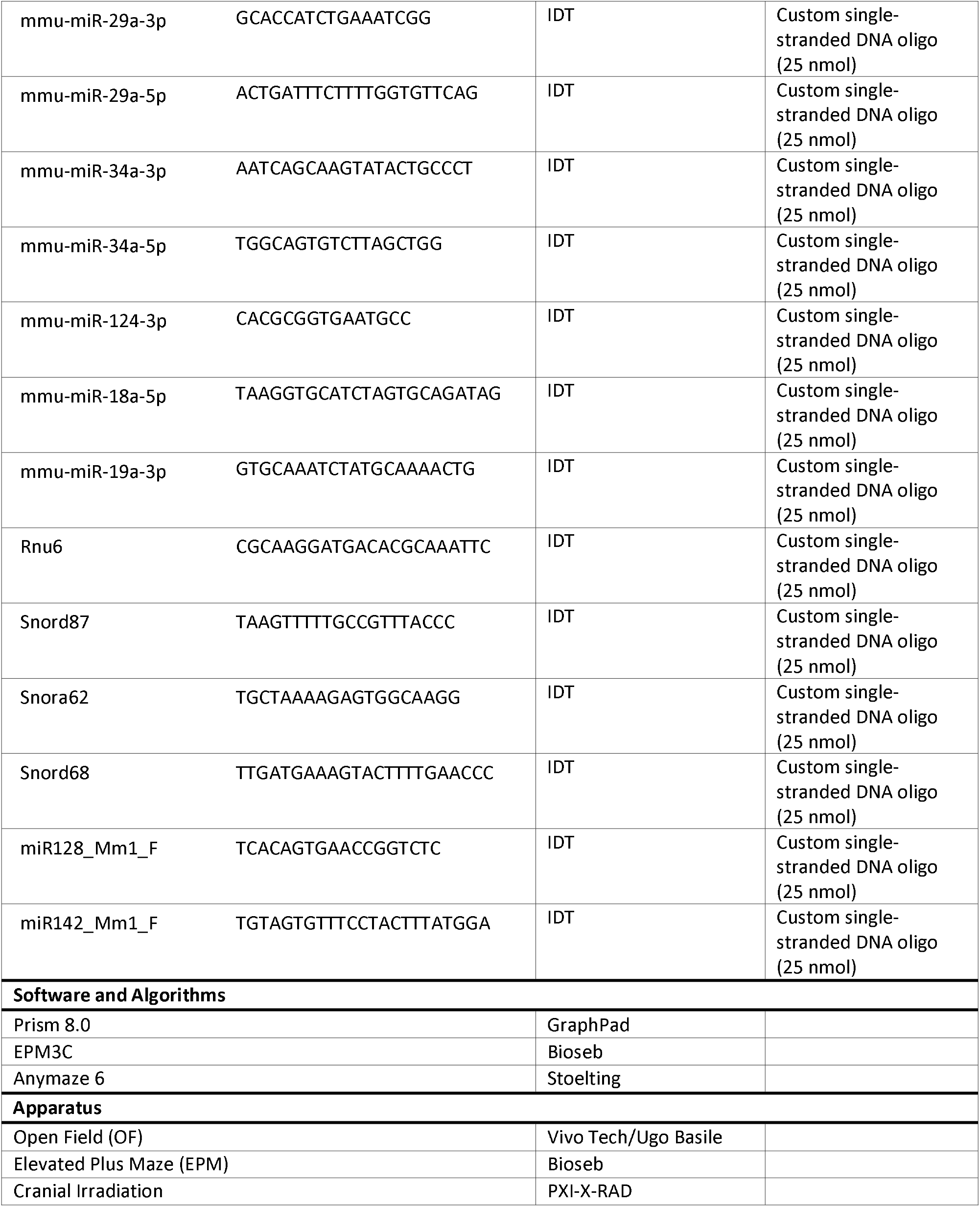

### CONTACT FOR REAGENT AND RESOURCE SHARING

Further information and requests for resources and reagents should be directed to and will be fulfilled by the lead contacts, Denis DAVID or René Hen.

### EXPERIMENTAL MODEL AND SUBJECT DETAILS

All mice were 7-8 weeks old, weighed 23-25g at the beginning of the treatment, and were maintained on a 12h light:12h dark schedule (lights on at 0600). They were housed in groups of five. Food and water were provided ad libitum. All testing was in conformity with the institutional guidelines that are in compliance with national and international laws and policies (Council directive # 87-848, October 19,1987, Ministère de ? Agriculture et de la Forêt, Service Vétérinaire de la Santé et de la Protection Animale; NIH Guide for the Care and Use of Laboratory Animal) and in compliance with protocols approved by the Institutional Animal Care and Use Committee (CEE26 authorization #4747-2016033116126273 at Université Paris-Saclay; Institutional Animal Care and Use Committee of Columbia University and the Research Foundation for Mental Hygiene, Inc.).

### METHODS DETAILS

#### Animals

##### Study in β-arrestin 2 KO mice

β-arrestin 2 +/− were provided from Bob Lefkowitz (Duke University). Male heterozygous β-Arr2 heterozygous female mutant β-Arr2 mice (age 4–6 months) were bred on a mixed S129/Sv × C57BL/6 genetic background at the University Paris-Saclay’s animal facility (Chatenay-Malabry, France). Resulting pups were genotyped by polymerase chain reaction (Beaulieu et al., 2008; David et al., 2009). Male heterozygous β-Arr2 Hets, β-Arr2 KO and their littermates (WT), 25–30 g body weight, were used to test the behavioral and neurogenic effects of a chronic treatment with fluoxetine (18 mg/kg/day) (See timeline of experiments in **Figure 1A**).

##### Study in CORT model

*Study in mice presenting an intact adult hippocampal neurogenesis*

Male C57BL/6JRj mice (Janvier Labs, France; 25–30 g bodyweight) were used to test the neurogenic effects of a chronic treatment with fluoxetine (18 mg/kg/day) in the CORT model (See timeline of experiments in **Figure 3A or Figure 6**).

##### Study in irradiated mice

Male C57BL/6Ntac mice (Taconic Farms, Germantown, NY, USA, 25–30 g body weight) were used to test the neurogenic effects of a chronic treatment with fluoxetine (18 mg/kg/day) in the CORT model after X-ray procedure (see timeline of experiments in **Figure 4A**).

##### Study in GFAP-TK+ mice

Transgenic GFAP-TK+ and transgenic GFAP-TK-mice were used to test the neurogenic effects of a chronic treatment with fluoxetine (18 mg/kg/day) in the CORT model after (Mendez-David et al., 2017; Schloesser et al., 2009; Snyder et al., 2011) (see timeline of experiments in **Figure 4C**). An agreement (license L-O 15-2015/0) between the NIH and the Université Paris-Saclay provides CESP/UMRS 1178 laboratory with the use of transgenic GFAP-TK mice, which were developed in the laboratory of Dr. Heather Cameron of the National Institute of Mental Health (NIMH).

##### Lipopolysaccharide study

Male C57BL/6JRj mice (Janvier Labs, France, 25-30 g body weight) locally injected in the dorsal dentate gyrus with Lipopolysaccharide were used to assess the role of inflammation on AHN (See timeline of experiments in **Figure 5A).**

#### Drugs and treatment

##### Study in β-arrestin 2 KO mice

Behavioral and neurogenic effects of a 4 week treatment with fluoxetine hydrochloride [Anawa Trading, Zurich, Switzerland, 160 mg/ml, equivalent to 18 mg/kg/day in drinking water, according (David et al., 2009)] were tested in male β-Arr2 Hets, β-Arr2 KO mice and their WT littermates **(Figure 1, 2, Supplemental Figure 2).** Similarly, neurogenic effects of a 4-week fluoxetine treatment were evaluated in male β-Arr1 KO and their littermates **(Supplemental Figure 1**).

##### Study in CORT model

Corticosterone [4-pregnen-11b-DIOL-3 2O-DIONE 21-hemi-succinate (35 μg/ml)] purchased from Sigma-Aldrich (Saint-Quentin Fallavier, France) was dissolved in a vehicle (0.45% hydroxypropyl-β-cyclodextrin (β-CD); Sigma-Aldrich). Fluoxetine hydrochloride (160 mg/ml, equivalent to 18 mg/kg/day) was purchased from Anawa Trading, (Zurich, Switzerland) and dissolved in 0.45% β-CD/corticosterone solution. Corticosterone-treated water was changed every 3 days to prevent any possible degradation as previously described (David et al., 2009; Mendez-David et al., 2014). Fluoxetine (18 mg/kg/day drinking water) was administered chronically to mice for 28 days, 4 weeks after chronic corticosterone and their effects on AHN were observed in mice presenting an intact **(Figures 3 and 6, Supplemental Figures 3)** or an ablation of **AHN (Figures 4, Supplemental Figures 4).**

##### Arresting Adult Hippocampal Neurogenesis

###### -X-irradiation

Male C57BL/6Ntac mice were anesthetized with ketamine and xylazine (75/20 mg/kg), placed in a stereotaxic frame, and exposed to cranial irradiation using a PXI X-RAD 320 X-ray system operated at 300 kV and 12 mA with a 2 mm Al filter according a well-described procedure (Anacker et al., 2018; Mendez-David et al., 2014). Animals were protected with a lead shield that covered the entire body, but left unshielded a 3.22 × 11-mm treatment field above the hippocampus (interaural 3.00 to 0.00) exposed to X-ray, thus effectively preventing irradiation from targeting the rest of the brain (Santarelli et al., 2003). The corrected dose rate was approximately 0.95 Gy per min at a source to skin distance of 36 cm. The procedure lasted 2 min and 39 sec, delivering a total of 2.5 Gy. Three 2.5 Gy doses were delivered on days 1, 4, and 7. This 7.5 Gy cumulative dose was determined from prior pilot experiments to be the minimum dosage necessary to result in permanent ablation of adult-born neurons in the dentate gyrus as assessed by expression of the immature neuronal marker doublecortin. The reason for using a fractionated paradigm rather than a single high dose of 7.5Gy is that the ablation is not permanent after a single high dose. Then, Sham or Irradiated animals were submitted to the CORT protocol in presence of fluoxetine (18 mg/kg/day in the drinking water during the last 4 weeks) and sacrificed 8 weeks after.

###### -Genetic ablation of adult hippocampal neurogenesis study

To arrest neurogenesis in GFAP-Tk+ Mice, ValGanciclovir (vGCV, Sigma-Aldrich, St Quentin Fallavier, France) – the L-valyl ester of ganciclovir - were given to GFAP-Tk+ mice and their littermates GFAP-Tk-from Monday to Friday during 12 weeks through the animals’ chow at a concentration of 165 mg/kg [SSNIFF, Soest, Germany, (Mendez-David et al., 2017)]. Then, 4 weeks after the start of vGCG protocol, GFAP-Tk+ animals and their littermates were submitted to the CORT protocol (35 μg/ml), in presence or absence of fluoxetine (18 mg/kg/ day in the drinking water) for the last 4 weeks of the protocol.

##### Lipopolysaccharide study

A week after receiving 5-Bromo-2-Deoxyuridine (BrdU), C57BL/6JRj mice were anesthetized with chloral hydrate (400 mg/kg, i.p.). Lipopolysaccharide from Escherichia coli 0111:B4 (LPS, Sigma-Aldrich, St Quentin-Fallavier, France) (1 μg) was infused perfused at a flow rate of 0.25 μL/min during 2 minutes (LEGATO™ 180 syringe pump, KD Scientific Inc., Holliston, MA, USA) in the dorsal dentate gyrus (stereotaxic coordinates in mm from bregma: A= −1.4, L= ± 1.0, V= −1.75; A, anterior; L, lateral; and V, ventral) (Chugh et al., 2013). Animals were sacrificed 24hrs, 3 or 7 days after LPS injection for immunochemistry.

#### Behavioral study in β-arrestin 2 −/− KO mice

The same cohort of animals was tested in three different behavioral models of anxiety and depression. Each animal, over a week, was successively tested in the Open Field (OF), Elevated Plus Maze (EPM), and Novelty Suppressed Feeding (NSF). Behavioral testing occurred during the light phase between 0700 and 1900. Behavioral paradigms occurred after 28 days of fluoxetine treatment **(Figure 1).**

##### Open Field (OF)

This test was performed as described by (Faye et al., 2019). Motor activity was quantified in four 39 × 39 cm Perspex plastic open field boxes (Vivo-tech/Ugo Basile, Salon de Provence, France). The apparatus was illuminated from the ground with special designed 40 x 40 cm Infra-red backlights (monochromatic wavelength 850 nm high homogeneity, Vivo-tech, Salon de Provence, France). Activity chambers were monitored by four black and white cameras with varifocal optics and polarizing filters (Vivo-tech, Salon de Provence, France). The whole set-up was controlled using ANYMAZE version 6 video tracking software (Stoelting Co/Vivo-tech, Salon de Provence, France). Dependent measures were time in the center over a 30 min for systemic administration or 6 min test period for optogenetic experiments, total ambulatory distance and ambulatory distance traveled in the center divided by total distance.

##### Elevated Plus Maze (EPM)

This test was performed as described by Faye and colleagues (Faye et al., 2019). The maze is a plus-cross-shaped apparatus, with two open arms and two arms closed by walls linked by a central platform 50 cm above the floor. Mice were individually put in the center of the maze facing an open arm and were allowed to explore the maze during 5 min. The time spent in and the number of entries into the open arms was used as an anxiety index. All parameters were measured using a videotracker (EPM3C, Bioseb, Vitrolles, France).

##### Novelty Suppressed Feeding (NSF)

The NSF is a conflict test that elicits competing motivations: the drive to eat and the fear of venturing into the center of a brightly lit arena. The latency to begin eating is used as an index of anxiety/depression-like behavior, because classical anxiolytic drugs as well as chronic antidepressants decrease this measure (David et al., 2009). The NSF test was carried out during a 15-min period as previously described. Briefly, the testing apparatus consisted of a plastic box (50×50×20 cm), the floor of which was covered with approximately 2 cm of wooden bedding. Twenty-four hours prior to behavioral testing, all food was removed from the home cage. At the time of testing, a single pellet of food (regular chow) was placed on a white paper platform positioned in the center of the box. Each animal was placed in a corner of the box, and a stopwatch was immediately started. The latency to eat (defined as the mouse sitting on its haunches and biting the pellet with the use of forepaws) was timed. Immediately afterwards, the animal was transferred to its home cage, and the amount of food consumed by the mouse in the subsequent 5 min was measured, serving as a control for change in appetite as a possible confounding factor.

##### Immunohistochemistry

The effects of chronic treatment with fluoxetine on AHN (proliferation and/or survival and/or maturation) were assessed in β-Arr1 KO, β-Arr2 Hets, β-Arr2 KO mice and their WT littermates, in the CORT-treated mice presenting an intact or ablation of AHN or infusion of LPS in the dorsal dentate gyrus.

Thus, at the end of each procedure, animals were anesthetized with ketamine and xylazine (100 mg/ml ketamine; 20 mg/ml xylazine), and then perfused transcardially (cold saline for 2 min, followed by 4% cold para-formaldehyde at 4 °C). The brains were then removed and cryoprotected in 30% sucrose and stored at 4 °C. Serial sections (35 μm) were cryosectioned through the entire hippocampus (−1.10 to −3.80 mm relative to Bregma according to Franklin and Paxino’s brain atlas (2008) (Franklin and Paxinos, 2007) stored in PBS with 0.1% NaN3.

Since LPS was only injected in the dorsal dentate gyrus, its consequences the contralateral side and also along the septotemporal axis of the dentate gyrus, were also analyzed. The coordinates used to dissociate the dorsal and ventral hippocampus were based on previous publications (Rainer et al., 2012): from −1.10 to −2.50 mm relative to Bregma for the dorsal hippocampus and from – 2.50 to −3.80 mm for the ventral hippocampus according to Franklin and Paxino’s brain atlas (Franklin and Paxinos, 2007).

##### Proliferation study

For Ki67 immunostaining, sections were incubated in 0.3% triton in PBS and 10% Normal Donkey Serum (NDS). Then sections were incubated overnight at 4°C with anti-rabbit Ki67 (from Vector; 1:100). After washing with PBS, sections were incubated for 2 h with secondary antibody (1:200 biotinylated donkey anti-rabbit). Cells were counted using a BX51 microscope (Olympus, Germany).

##### Survival study

Mice were administered with 5-Bromo-2-Deoxyuridine (BrdU) (150 mg/kg, twice a day during 3 days) before the start of the different treatments and processed as described by (Mendez-David et al., 2014). BrdU positive cells were counted using a BX51 microscope (Olympus, Germany).

##### Doublecortin (DCX) Labelling

The immunohistochemistry protocol was adapted from (David et al., 2009). DCX-positive (DCX+) cells were subcategorized according to their dendritic morphology: DCX+ cells with no tertiary dendritic processes and DCX+ cells with tertiary (or higher order) dendrites.

##### Calretinin (CR) labelling

For CR immunostaining, sections were incubated in 0.3% triton in PBS and 10% Normal Donkey Serum (NDS). Then sections were incubated overnight at 4°C with anti-rabbit CR polyclonal antibody (from Swant; 1:10,000). After washing with PBS, sections were incubated for 2 h with secondary antibody (1:500 Jackson ImmunoResearch). Cells were counted using a BX51 microscope (Olympus, Germany).

##### Immunohistochemistry and confocal imaging for maturation study

Immunohistochemistry was performed in the following steps: 2 h incubation in 1:1 formamide/2xSSC at 65 °C, 5 min rinse in 2xSSC, 30 min incubation in 2 N HCl at 37 °C, and 10 min rinse in 0.1 M boric acid, pH 8.5, 2 h incubation in 0.1 M PBS with 0.3% Triton X-100, and 5% normal donkey serum. Sections were then incubated overnight at 4 °C in primary antibodies for doublecortin (goat 1:500; Santa Cruz Biotechnology, Santa Cruz, CA), bromodeoxyuridine (BrdU; rat; 1:100; Serotec, Oxford, UK) and neuronal-specific nuclear protein (NeuN) (mouse; 1:500; Chemicon, Temecula, CA). Then fluorescent secondary antibodies were used. All secondary antibodies were purchased from Jackson ImmunoResearch (1:500). Approximately 6 sections per animal and 20-30 BrdU+ cells per treatment group were analysed (n = 4-5 animals per conditions). To identify colocalization of BrdU+/DCX+/NeuN+, z-stacks of immunolabeled sections were obtained using a confocal scanning microscope (Leica TCS SP8, Leica Microsystems Inc.) equipped with three simultaneous PMT detectors.

#### Hippocampal doublecortin and MicroRNA expression

##### Hippocampal microdissection and preparation of total RNA

Mouse hippocampal tissues were isolated and frozen in liquid nitrogen, and tissue was stored at −80 °C. Total RNA was isolated from frozen tissue samples using TRIzol (Invitrogen, Carlsbad, Calif.) and according to the manufacturer’s protocol. RNA quality was assessed using using a BioMate™3S spectrophotometer (Thermo Scientific). RNA quality was assessed using the Biophotometer (Eppendorf) and gel electrophoresis with the RNA LabChip^®^ 6000 Nano kit (Bioanalyzer^®^ 2000, Agilent Technologies). RNA integrity was evaluated by capillary electrophoresis using RNA 6000 Nano chips and the Bioanalyzer 2100 (Agilent Technologies). All RIN score values were above 8.0.

##### RT-qPCR analysis

For quantification of mRNA or miRNA expression, first strand cDNA was synthesized from 0.2 to 1 μg of total RNA, using the miSCRIPT^®^ II RT kit (Qiagen), according to the manufacturer’s instructions. PCR forward primer specific to each target gene was designed from the RefSeq sequence using Primer3Plus software (https://primer3plus.com/cgi-bin/dev/primer3plus.cgi). The cDNA synthesized from 3 ng of total RNA was amplified in a CFX384™ real time thermal cycler (Bio-Rad), either for miR quantitation using the miSCRIPT SYBR Green PCR kit (Qiagen) according to manufacturer’s instructions, with final concentrations of 500-nM forward primer and 1X reverse universal primer, in triplicate 10-μl reactions, by 40 cycles (94°C 10 s; 56°C 25 s; 70°C 20 s), or for mRNA quantitation using the Sso Advanced SYBR Green PCR kit (Bio-Rad) according to manufacturer’s instructions, with 500-nM final concentrations of each primer, in triplicate 10-μl reactions, by 45 two-step cycles (94°C 10 s; 60°C 25 s). ‘No RT’ controls were amplified to check for the absence of for genomic DNA contamination, and melting curve analysis was performed to assess the purity of the PCR products. PCR efficiencies calculated for each gene from the slopes of calibration curves generated from the pool of all cDNA samples were above 95%. Gapdh and Actb genes were used as references for normalization of mRNA expression results, and Rnu6, Snora62, Snord68 and Snord87 for miR expression results. The normalized relative expression of target genes in samples was determined using the ^ΔΔ^Cq method with correction for PCR efficiencies, where *NRQ* 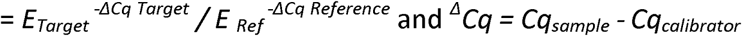 (Hellemans et al., 2007). Final results were expressed as the n-fold differences in target gene expression in treated vs vehicle cells. Mean values ± SEM from both mice groups are presented.

#### Statistical analysis

Results from data analyses, expressed as mean ± SEM were analyzed using Prism 8.3 software (Graphpad, San Diego, CA, USA). For all experiments, unpaired t-test, one-way ANOVA with repeated measures or two-way ANOVAs were applied to the data as appropriate. Significant main effects and/or interactions were followed by Fisher’s PLSD post-hoc analysis. Statistical significance was set at p<0.05. All statistical tests and p values are listed in Supplemental Tables (Supplemental Tables 1-9).

## REFERENCES

Anacker, C., Luna, V.M., Stevens, G.S., Millette, A., Shores, R., Jimenez, J.C., Chen, B., and Hen, R. (2018). Hippocampal neurogenesis confers stress resilience by inhibiting the ventral dentate gyrus. Nature 559, 98–102.

Asth, L., Ruzza, C., Malfacini, D., Medeiros, I., Guerrini, R., Zaveri, N.T., Gavioli, E.C., and Calo, G. (2016). Beta-arrestin 2 rather than G protein efficacy determines the anxiolytic-versus antidepressant-like effects of nociceptin/orphanin FQ receptor ligands. Neuropharmacology 105, 434–442.

Baptista, P., and Andrade, J.P. (2018). Adult Hippocampal Neurogenesis: Regulation and Possible Functional and Clinical Correlates. Front Neuroanat 12, 44.

Beaulieu, J.M., Marion, S., Rodriguiz, R.M., Medvedev, I.O., Sotnikova, T.D., Ghisi, V., Wetsel, W.C., Lefkowitz, R.J., Gainetdinov, R.R., and Caron, M.G. (2008). A beta-arrestin 2 signaling complex mediates lithium action on behavior. Cell 132, 125–136.

Bohn, L.M., and Schmid, C.L. (2010). Serotonin receptor signaling and regulation via beta-arrestins. Crit Rev Biochem Mol Biol 45, 555–566.

Boldrini, M., Fulmore, C.A., Tartt, A.N., Simeon, L.R., Pavlova, I., Poposka, V., Rosoklija, G.B., Stankov, A., Arango, V., Dwork, A.J., et al. (2018). Human Hippocampal Neurogenesis Persists throughout Aging. Cell Stem Cell 22, 589–599 e585.

Boldrini, M., Galfalvy, H., Dwork, A.J., Rosoklija, G.B., Trencevska-Ivanovska, I., Pavlovski, G., Hen, R., Arango, V., and Mann, J.J. (2019a). Resilience Is Associated With Larger Dentate Gyrus, While Suicide Decedents With Major Depressive Disorder Have Fewer Granule Neurons. Biol Psychiatry 85, 850–862.

Boldrini, M., Galfalvy, H., Dwork, A.J., Rosoklija, G.B., Trencevska-Ivanovska, I., Pavlovski, G., Hen, R., Arango, V., and Mann, J.J. (2019b). Resilience Is Associated With Larger Dentate Gyrus, While Suicide Decedents With Major Depressive Disorder Have Fewer Granule Neurons. Biol Psychiatry.

Burghardt, N.S., Park, E.H., Hen, R., and Fenton, A.A. (2012). Adult-born hippocampal neurons promote cognitive flexibility in mice. Hippocampus 22, 1795–1808.

Chugh, D., Nilsson, P., Afjei, S.A., Bakochi, A., and Ekdahl, C.T. (2013). Brain inflammation induces post-synaptic changes during early synapse formation in adult-born hippocampal neurons. Exp Neurol 250, 176–188.

David, D.J., Samuels, B.A., Rainer, Q., Wang, J.W., Marsteller, D., Mendez, I., Drew, M., Craig, D.A., Guiard, B.P., Guilloux, J.P., et al. (2009). Neurogenesis-dependent and -independent effects of fluoxetine in an animal model of anxiety/depression. Neuron 62, 479–493.

Denny, C.A., Burghardt, N.S., Schachter, D.M., Hen, R., and Drew, M.R. (2012). 4-to 6-week-old adult-born hippocampal neurons influence novelty-evoked exploration and contextual fear conditioning. Hippocampus 22, 1188–1201.

Dhaliwal, J., Xi, Y., Bruel-Jungerman, E., Germain, J., Francis, F., and Lagace, D.C. (2015). Doublecortin (DCX) is not Essential for Survival and Differentiation of Newborn Neurons in the Adult Mouse Dentate Gyrus. Front Neurosci 9, 494.

Diaz, S.L., Doly, S., Narboux-Neme, N., Fernandez, S., Mazot, P., Banas, S.M., Boutourlinsky, K., Moutkine, I., Belmer, A., Roumier, A., et al. (2012). 5-HT(2B) receptors are required for serotonin-selective antidepressant actions. Mol Psychiatry 17, 154–163.

Dwivedi, Y. (2013). microRNAs as Biomarker in Depression Pathogenesis. Ann Psychiatry Ment Health 1, 1003.

Dwivedi, Y. (2014). Emerging role of microRNAs in major depressive disorder: diagnosis and therapeutic implications. Dialogues Clin Neurosci 16, 43–61.

Encinas, J.M., Vaahtokari, A., and Enikolopov, G. (2006). Fluoxetine targets early progenitor cells in the adult brain. Proc Natl Acad Sci U S A 103, 8233–8238.

Evangelisti, C., Florian, M.C., Massimi, I., Dominici, C., Giannini, G., Galardi, S., Bue, M.C., Massalini, S., McDowell, H.P., Messi, E., et al. (2009). MiR-128 up-regulation inhibits Reelin and DCX expression and reduces neuroblastoma cell motility and invasiveness. FASEB J 23, 4276–4287.

Faye, C., Hen, R., Guiard, B.P., Denny, C.A., Gardier, A.M., Mendez-David, I., and David, D.J. (2019). Rapid Anxiolytic Effects of RS67333, a Serotonin Type 4 Receptor Agonist, and Diazepam, a Benzodiazepine, Are Mediated by Projections From the Prefrontal Cortex to the Dorsal Raphe Nucleus. Biol Psychiatry.

Flor-Garcia, M., Terreros-Roncal, J., Moreno-Jimenez, E.P., Avila, J., Rabano, A., and Llorens-Martin, M. (2020). Unraveling human adult hippocampal neurogenesis. Nat Protoc.

Franklin, K.B.J., and Paxinos, G. (2007). The Mouse Brain in stereotaxic coordinates. Academic Press 3rd Edition.

Hellemans, J., Mortier, G., De Paepe, A., Speleman, F., and Vandesompele, J. (2007). qBase relative quantification framework and software for management and automated analysis of real-time quantitative PCR data. Genome Biol 8, R19.

Jin, J., Kim, S.N., Liu, X., Zhang, H., Zhang, C., Seo, J.S., Kim, Y., and Sun, T. (2016). miR-17-92 Cluster Regulates Adult Hippocampal Neurogenesis, Anxiety, and Depression. Cell Rep 16, 1653–1663.

Kempermann, G., Song, H., and Gage, F.H. (2015). Neurogenesis in the Adult Hippocampus. Cold Spring Harb Perspect Biol 7, a018812.

Klempin, F., Kronenberg, G., Cheung, G., Kettenmann, H., and Kempermann, G. (2011). Properties of doublecortin-(DCX)-expressing cells in the piriform cortex compared to the neurogenic dentate gyrus of adult mice. PLoS One 6, e25760.

Kobayashi, K., Ikeda, Y., Sakai, A., Yamasaki, N., Haneda, E., Miyakawa, T., and Suzuki, H. (2010). Reversal of hippocampal neuronal maturation by serotonergic antidepressants. Proc Natl Acad Sci U S A 107, 8434–8439.

Kohler, S.J., Williams, N.I., Stanton, G.B., Cameron, J.L., and Greenough, W.T. (2011). Maturation time of new granule cells in the dentate gyrus of adult macaque monkeys exceeds six months. Proc Natl Acad Sci U S A 108, 10326–10331.

Kremer, T., Jagasia, R., Herrmann, A., Matile, H., Borroni, E., Francis, F., Kuhn, H.G., and Czech, C. (2013). Analysis of adult neurogenesis: evidence for a prominent “non-neurogenic” DCX-protein pool in rodent brain. PLoS One 8, e59269.

Lino de Oliveira, C., Bolzan, J.A., Surget, A., and Belzung, C. (2020). Do antidepressants promote neurogenesis in adult hippocampus? A systematic review and meta-analysis on naive rodents. Pharmacol Ther, 107515.

Ma, Z., Zang, T., Birnbaum, S.G., Wang, Z., Johnson, J.E., Zhang, C.L., and Parada, L.F. (2017). TrkB dependent adult hippocampal progenitor differentiation mediates sustained ketamine antidepressant response. Nat Commun 8, 1668.

Malberg, J.E., Eisch, A.J., Nestler, E.J., and Duman, R.S. (2000). Chronic antidepressant treatment increases neurogenesis in adult rat hippocampus. J Neurosci 20, 9104–9110.

Mendez-David, I., David, D.J., Darcet, F., Wu, M.V., Kerdine-Romer, S., Gardier, A.M., and Hen, R. (2014). Rapid anxiolytic effects of a 5-HT(4) receptor agonist are mediated by a neurogenesis-independent mechanism. Neuropsychopharmacology 39, 1366–1378.

Mendez-David, I., Guilloux, J.P., Papp, M., Tritschler, L., Mocaer, E., Gardier, A.M., Bretin, S., and David, D.J. (2017). S 47445 Produces Antidepressant- and Anxiolytic-Like Effects through Neurogenesis Dependent and Independent Mechanisms. Front Pharmacol 8, 462.

Moreno-Jimenez, E.P., Flor-Garcia, M., Terreros-Roncal, J., Rabano, A., Cafini, F., Pallas-Bazarra, N., Avila, J., and Llorens-Martin, M. (2019). Adult hippocampal neurogenesis is abundant in neurologically healthy subjects and drops sharply in patients with Alzheimer’s disease. Nat Med 25, 554–560.

Ngwenya, L.B., Heyworth, N.C., Shwe, Y., Moore, T.L., and Rosene, D.L. (2015). Age-related changes in dentate gyrus cell numbers, neurogenesis, and associations with cognitive impairments in the rhesus monkey. Front Syst Neurosci 9, 102.

Ohira, K., Hagihara, H., Miwa, M., Nakamura, K., and Miyakawa, T. (2019). Fluoxetine-induced dematuration of hippocampal neurons and adult cortical neurogenesis in the common marmoset. Mol Brain 12, 69.

Opal, M.D., Klenotich, S.C., Morais, M., Bessa, J., Winkle, J., Doukas, D., Kay, L.J., Sousa, N., and Dulawa, S.M. (2014). Serotonin 2C receptor antagonists induce fast-onset antidepressant effects. Mol Psychiatry 19, 1106–1114.

Plumpe, T., Ehninger, D., Steiner, B., Klempin, F., Jessberger, S., Brandt, M., Romer, B., Rodríguez, G.R., Kronenberg, G., and Kempermann, G. (2006). Variability of doublecortin-associated dendrite maturation in adult hippocampal neurogenesis is independent of the regulation of precursor cell proliferation. BMC Neurosci 7, 77.

Pytka, K., Gluch-Lutwin, M., Zmudzka, E., Salaciak, K., Siwek, A., Niemczyk, K., Walczak, M., Smolik, M., Olczyk, A., Galuszka, A., et al. (2018). HBK-17, a 5-HT1A Receptor Ligand With Anxiolytic-Like Activity, Preferentially Activates ss-Arrestin Signaling. Front Pharmacol 9, 1146.

Rainer, Q., Xia, L., Guilloux, J.P., Gabriel, C., Mocaer, E., Hen, R., Enhamre, E., Gardier, A.M., and David, D.J. (2012). Beneficial behavioural and neurogenic effects of agomelatine in a model of depression/anxiety. Int J Neuropsychopharmacol 15, 321–335.

Robinson, S.A., Brookshire, B.R., and Lucki, I. (2016). Corticosterone exposure augments sensitivity to the behavioral and neuroplastic effects of fluoxetine in C57BL/6 mice. Neurobiol Stress 3, 34–42.

Saaber, F., Schutz, D., Miess, E., Abe, P., Desikan, S., Ashok Kumar, P., Balk, S., Huang, K., Beaulieu, J.M., Schulz, S., et al. (2019). ACKR3 Regulation of Neuronal Migration Requires ACKR3 Phosphorylation, but Not beta-Arrestin. Cell Rep 26, 1473–1488 e1479.

Sagi, Y., Medrihan, L., George, K., Barney, M., McCabe, K.A., and Greengard, P. (2019). Emergence of 5-HT5A signaling in parvalbumin neurons mediates delayed antidepressant action. Mol Psychiatry.

Sahay, A., Scobie, K.N., Hill, A.S., O’Carroll, C.M., Kheirbek, M.A., Burghardt, N.S., Fenton, A.A., Dranovsky, A., and Hen, R. (2011). Increasing adult hippocampal neurogenesis is sufficient to improve pattern separation. Nature 472, 466–470.

Samuels, B.A., Mendez-David, I., Faye, C., David, S.A., Pierz, K.A., Gardier, A.M., Hen, R., and David, D.J. (2016). Serotonin 1A and Serotonin 4 Receptors: Essential Mediators of the Neurogenic and Behavioral Actions of Antidepressants. Neuroscientist 22, 26–45.

Sanai, N., Nguyen, T., Ihrie, R.A., Mirzadeh, Z., Tsai, H.H., Wong, M., Gupta, N., Berger, M.S., Huang, E., Garcia-Verdugo, J.M., et al. (2011). Corridors of migrating neurons in the human brain and their decline during infancy. Nature 478, 382–386.

Santarelli, L., Saxe, M., Gross, C., Surget, A., Battaglia, F., Dulawa, S., Weisstaub, N., Lee, J., Duman, R., Arancio, O., et al. (2003). Requirement of hippocampal neurogenesis for the behavioral effects of antidepressants. Science 301, 805–809.

Saxe, M.D., Battaglia, F., Wang, J.W., Malleret, G., David, D.J., Monckton, J.E., Garcia, A.D., Sofroniew, M.V., Kandel, E.R., Santarelli, L., et al. (2006). Ablation of hippocampal neurogenesis impairs contextual fear conditioning and synaptic plasticity in the dentate gyrus. Proc Natl Acad Sci USA 103, 17501–17506.

Schloesser, R.J., Manji, H.K., and Martinowich, K. (2009). Suppression of adult neurogenesis leads to an increased hypothalamo-pituitary-adrenal axis response. Neuroreport 20, 553–557.

Snyder, J.S. (2019). Recalibrating the Relevance of Adult Neurogenesis. Trends Neurosci 42, 164–178.

Snyder, J.S., Grigereit, L., Russo, A., Seib, D.R., Brewer, M., Pickel, J., and Cameron, H.A. (2016). A Transgenic Rat for Specifically Inhibiting Adult Neurogenesis. eNeuro 3.

Snyder, J.S., Soumier, A., Brewer, M., Pickel, J., and Cameron, H.A. (2011). Adult hippocampal neurogenesis buffers stress responses and depressive behaviour. Nature 476, 458–461.

Sorrells, S.F., Paredes, M.F., Cebrian-Silla, A., Sandoval, K., Qi, D., Kelley, K.W., James, D., Mayer, S., Chang, J., Auguste, K.I., et al. (2018). Human hippocampal neurogenesis drops sharply in children to undetectable levels in adults. Nature 555, 377–381.

Spalding, K.L., Bergmann, O., Alkass, K., Bernard, S., Salehpour, M., Huttner, H.B., Bostrom, E., Westerlund, I., Vial, C., Buchholz, B.A., et al. (2013). Dynamics of hippocampal neurogenesis in adult humans. Cell 153, 1219–1227.

Svenningsson, P., Chergui, K., Rachleff, I., Flajolet, M., Zhang, X., El Yacoubi, M., Vaugeois, J.M., Nomikos, G.G., and Greengard, P. (2006). Alterations in 5-HT1B receptor function by p11 in depression-like states. Science 311, 77–80.

Tartt, A.N., Fulmore, C.A., Liu, Y., Rosoklija, G.B., Dwork, A.J., Arango, V., Hen, R., Mann, J.J., and Boldrini, M. (2018). Considerations for Assessing the Extent of Hippocampal Neurogenesis in the Adult and Aging Human Brain. Cell Stem Cell 23, 782–783.

Toda, T., Parylak, S.L., Linker, S.B., and Gage, F.H. (2019). The role of adult hippocampal neurogenesis in brain health and disease. Mol Psychiatry 24, 67–87.

Verwer, R.W., Sluiter, A.A., Balesar, R.A., Baayen, J.C., Noske, D.P., Dirven, C.M., Wouda, J., van Dam, A.M., Lucassen, P.J., and Swaab, D.F. (2007). Mature astrocytes in the adult human neocortex express the early neuronal marker doublecortin. Brain 130, 3321–3335.

Wang, J.W., David, D.J., Monckton, J.E., Battaglia, F., and Hen, R. (2008). Chronic fluoxetine stimulates maturation and synaptic plasticity of adult-born hippocampal granule cells. J Neurosci 28, 1374–1384.

