## Supplemental Data for "A non-linear relation between levels of adult hippocampal neurogenesis and expression of the immature neuron marker doublecortin"

______________________________________________________________________________

**SUPPLEMENTAL FIGURES**

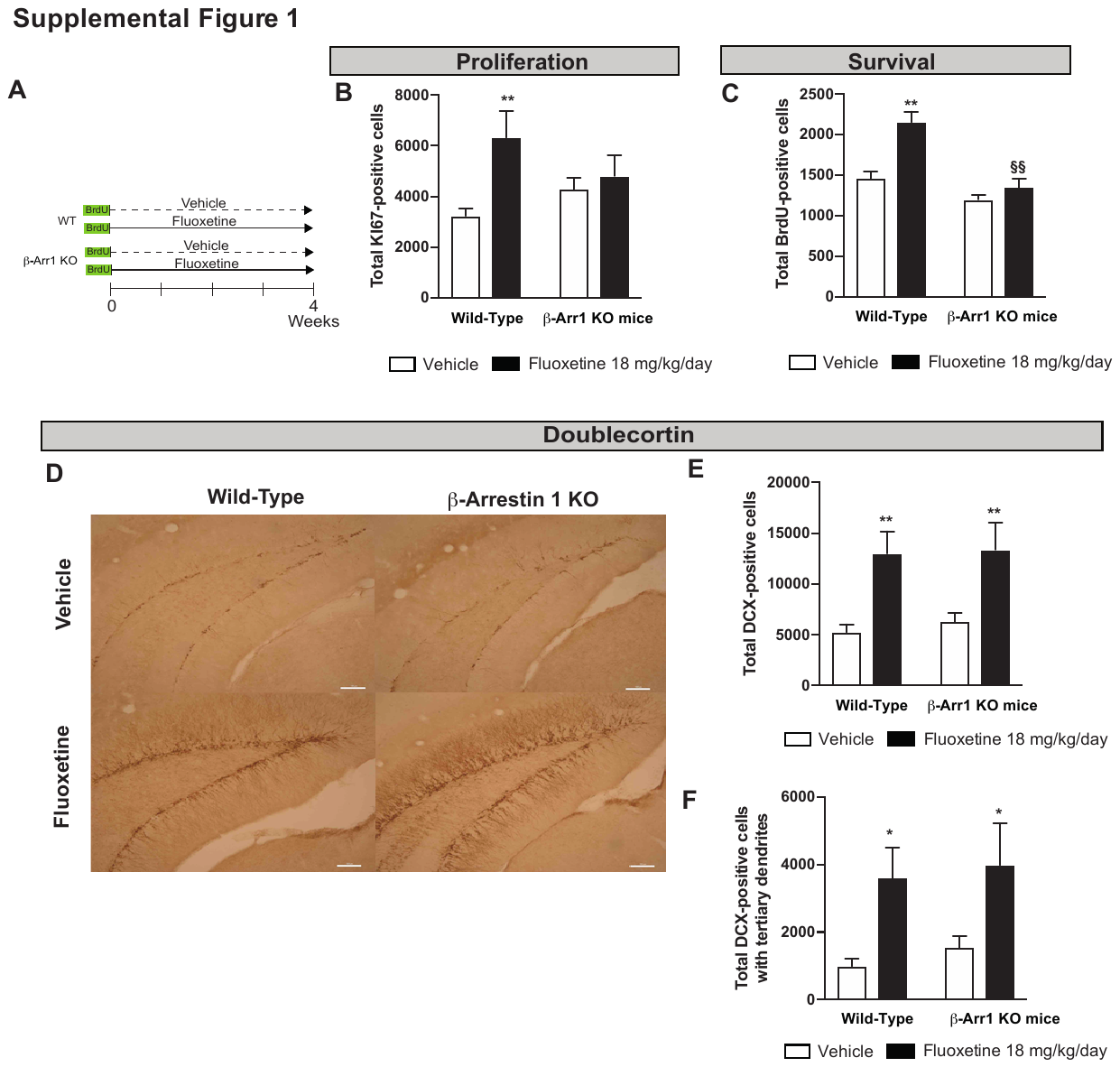

**Supplemental Figure 1: β-arrestin-1 expression is required for the effect of chronic fluoxetine on proliferation and survival of newborn cells but not maturation of young neurons.**

Experimental protocol timeline (A) to assess neurogenic consequences of a 4-week fluoxetine treatment (18 mg/kg/day in the drinking water) in β-arrestin-1 knockout mice (KO) mice compared with their Wild-Types littermates (WT). To evaluate survival of newborn cells, 5-Bromo-2-Deoxyuridine (BrdU) was administered twice a day at 150 mg/kg during 3 days, 4 weeks before sacrifice.

The effects of 28 days of treatment with fluoxetine (18 mg/kg/day) on cell proliferation (B), cell survival (C) and maturation of young neurons (C-F) were compare to those of vehicle in β-arrestin-1 knockout mice (KO) mice compared with their Wild-Types littermates (WT).

Images of doublecortin staining following vehicle or fluoxetine treatment in β-arrestin-1 KO mice and their WT littermates (10x magnification, scale bar= 100 μm) (C, F).

Proliferation (B) and survival (C) are measured as Ki67+ and BrdU+ cells respectively. Maturation was characterized by the total number of DCX+ cells (E), the number of DCX+ cells with tertiary dendrites (F).

Values plot are mean ± SEM (n = 3–7 animals/group). Data were analyzed with a two-way ANOVA (see supplementary table 3). **p< 0.01 for comparisons to the appropriate genotype/vehicle-treated group; #p< 0.05 for comparisons to WT/vehicle-treated group; §§p< 0.01 for comparisons to the WT/fluoxetine-treated group (n= 7/9 animals /group).

**Supplemental Figure 2: β-arrestin-2 modulates doublecortin expression in the piriform cortex of chronic fluoxetine-treated mice.**

Images of doublecortin staining following vehicle or fluoxetine (18 mg/kg/day) treatment in β-arrestin-2 KO mice and their WT littermates (4x magnification, scale bar= 100 μm and insert 10x magnification, scale bar= 100 μm) (A).

Maturation was characterized by the total number of DCX+ cells (B), the number of DCX+ cells with tertiary dendrites (C).

Values plot are mean ± SEM (n = 3–5 animals/group). Data were analyzed with a two-way ANOVA (see supplementary table 4).*p< 0.05; **p< 0.01 for comparisons to the appropriate genotype/vehicle-treated group; #p< 0.05 for comparisons to WT/vehicle-treated group; §p<0.05, §§p< 0.01 comparisons to the WT/fluoxetine-treated group.

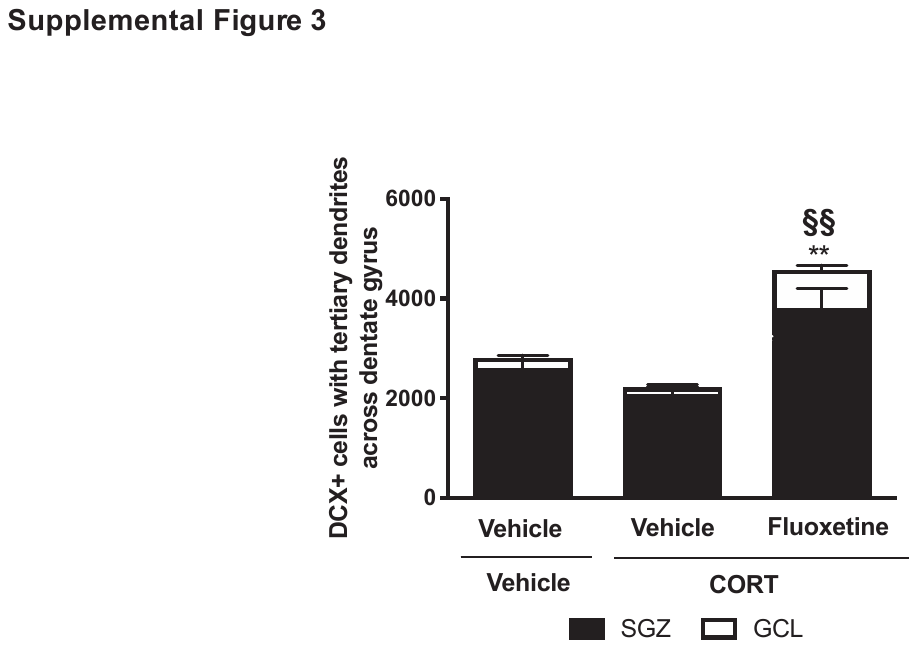

**Supplemental Figure 3: Chronic fluoxetine increases the number of doublecortin-positive cells with tertiary dendrites not only in the SGZ but also inside the granule cell layer (GCL).**

Male C57BL/6JRj mice (Janvier Labs, France; 25–30 g body weight) were submitted to the CORT protocol in presence of fluoxetine (18 mg/kg/day in the drinking water during the last 4 weeks) and sacrificed 8 weeks after. The effects of chronic fluoxetine (18 mg/kg/day) in corticosterone -treated animals on the number of DCX+ cells with tertiary dendrites were measured in the subgranular zone (SGZ) and the granule cell layer (GCL).

Values plot are mean ± SEM (n = 3 animals/group). Data were analyzed with a two-way ANOVA (see supplementary table 6). Significant main effects and/or interactions were followed by Fisher’s PLSD post-hoc analysis. **p< 0.01 for comparisons between fluoxetine-treated groups and vehicle-treated groups in CORT mice in SGZ or GCL; §§p< 0.01 comparisons between fluoxetine-treated CORT groups and vehicle treated control groups in SGZ or GCL.

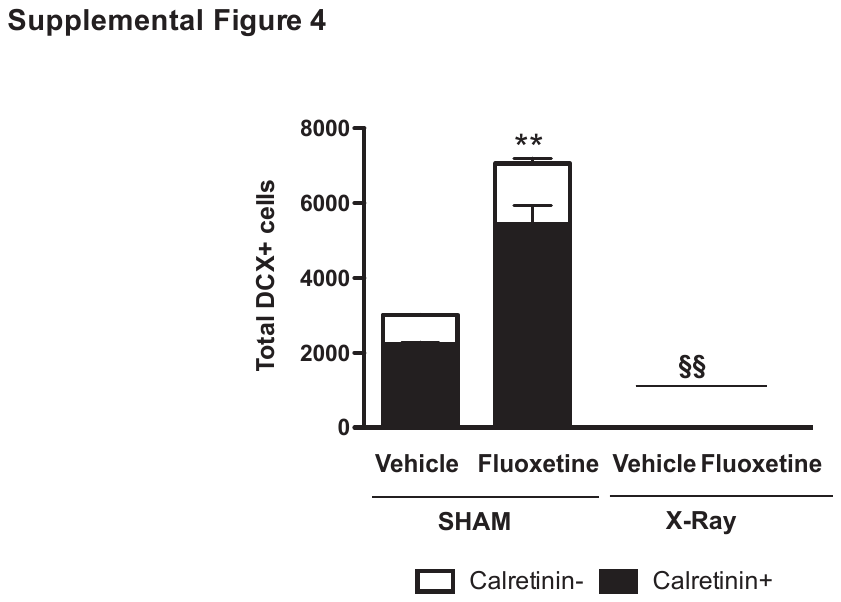

**Supplemental Figure 4: A majority of doublecortin-positive cells co-expressed calretinin and both populations disappear after X-irradiation.**

Sham or Irradiated animals (X-Ray) were submitted to the CORT protocol in presence of fluoxetine (18 mg/kg/day in the drinking water during the last 4 weeks) and sacrificed 8 weeks after. We counted cells expressing doublecortin as well as cells expressing both doublecortin and calretinin after fluoxetine treatment in sham and X-irradiated animals. Values are mean ± SEM (n = 5 animals/group). Data were analyzed with a two-way ANOVA (see supplementary table 7). Significant main effects and/or interactions were followed by Fisher’s PLSD post-hoc analysis **p< 0.01 for comparisons between fluoxetine and vehicle-treated CORT group. §§p< 0.01 for comparisons between sham and irradiated-treated groups.

**SUPPLEMENTAL TABLES**

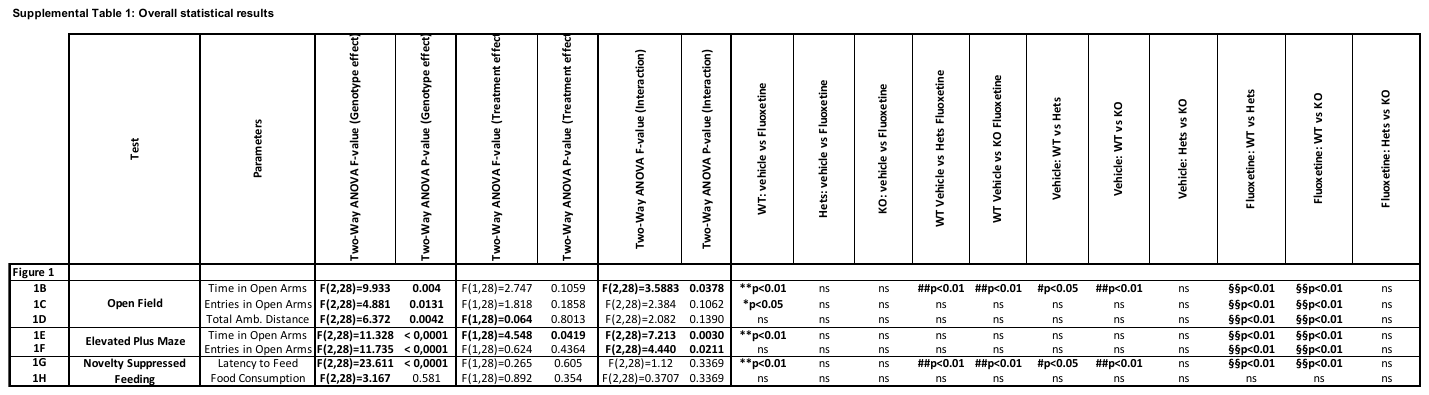

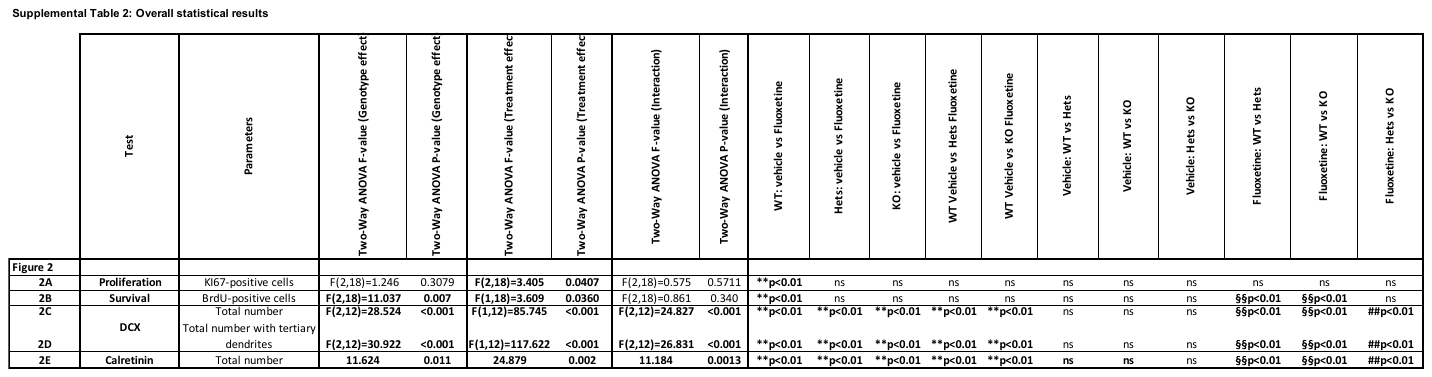

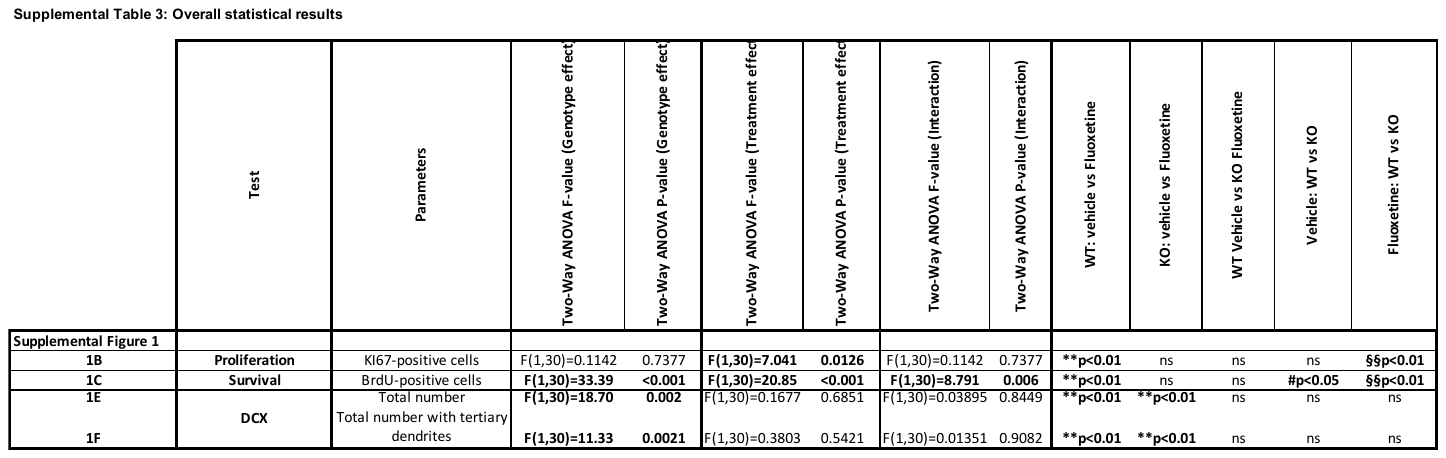

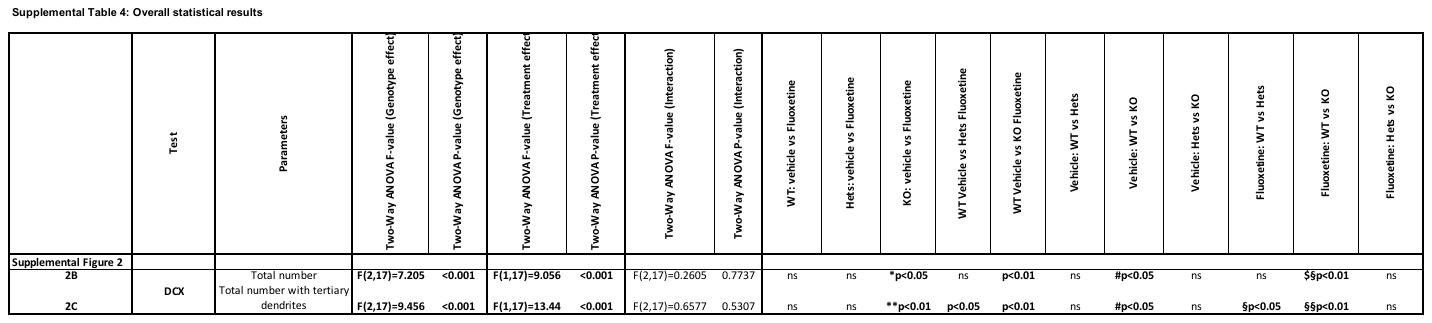

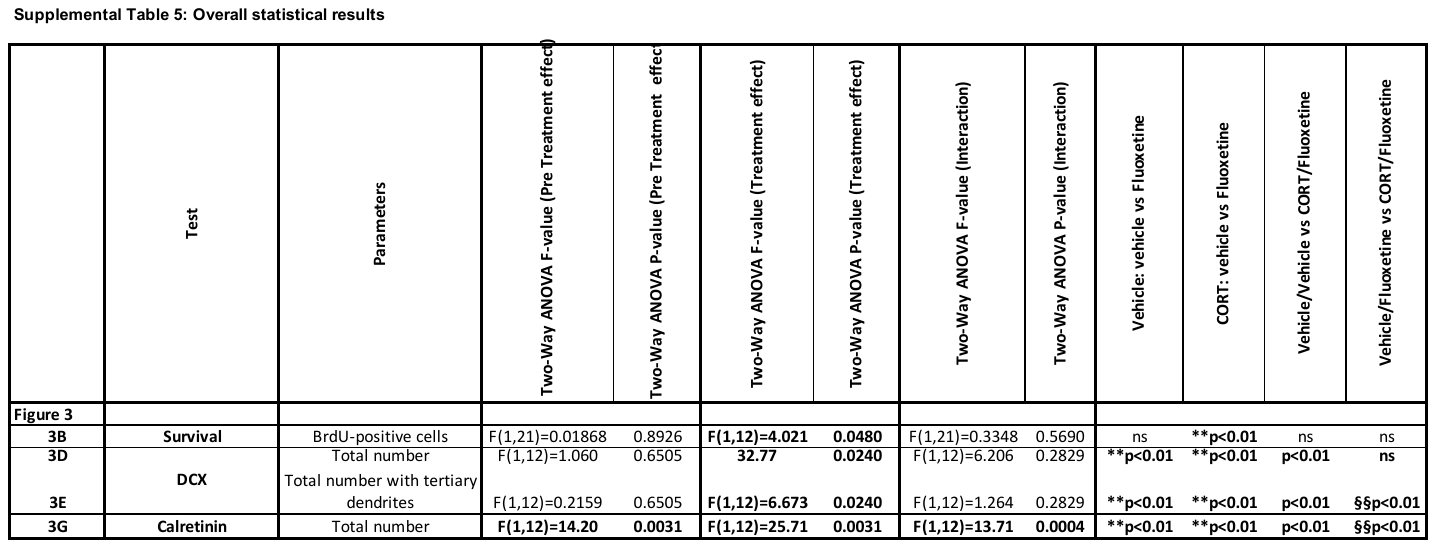

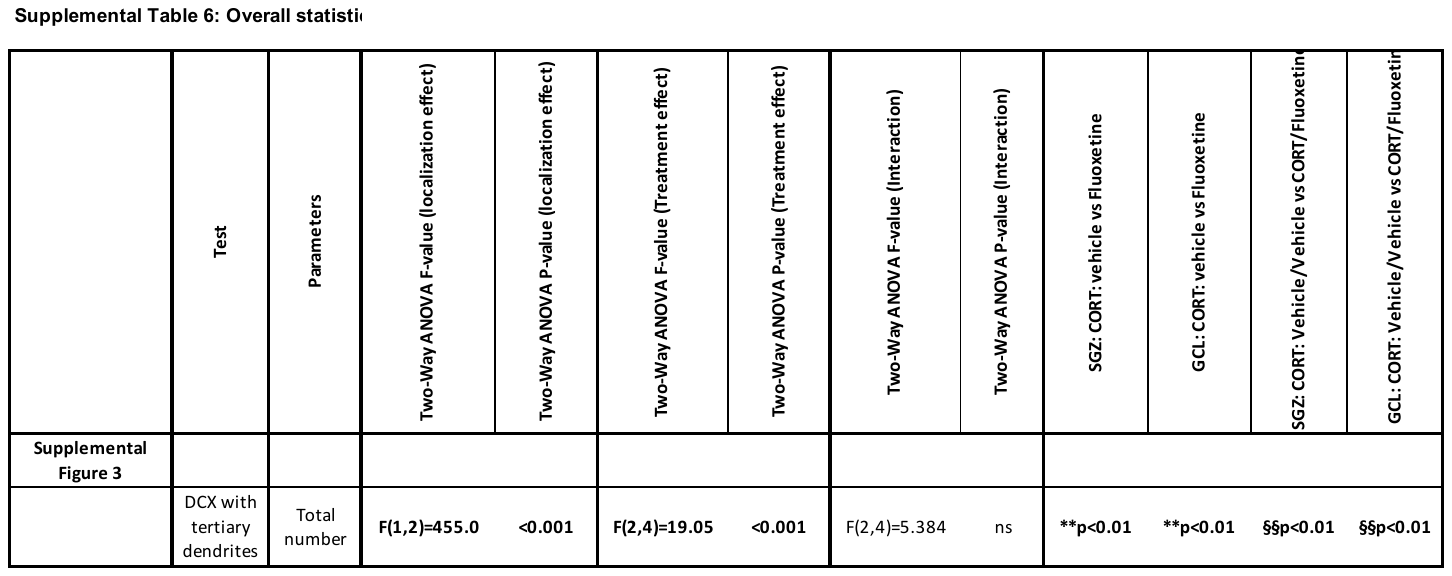

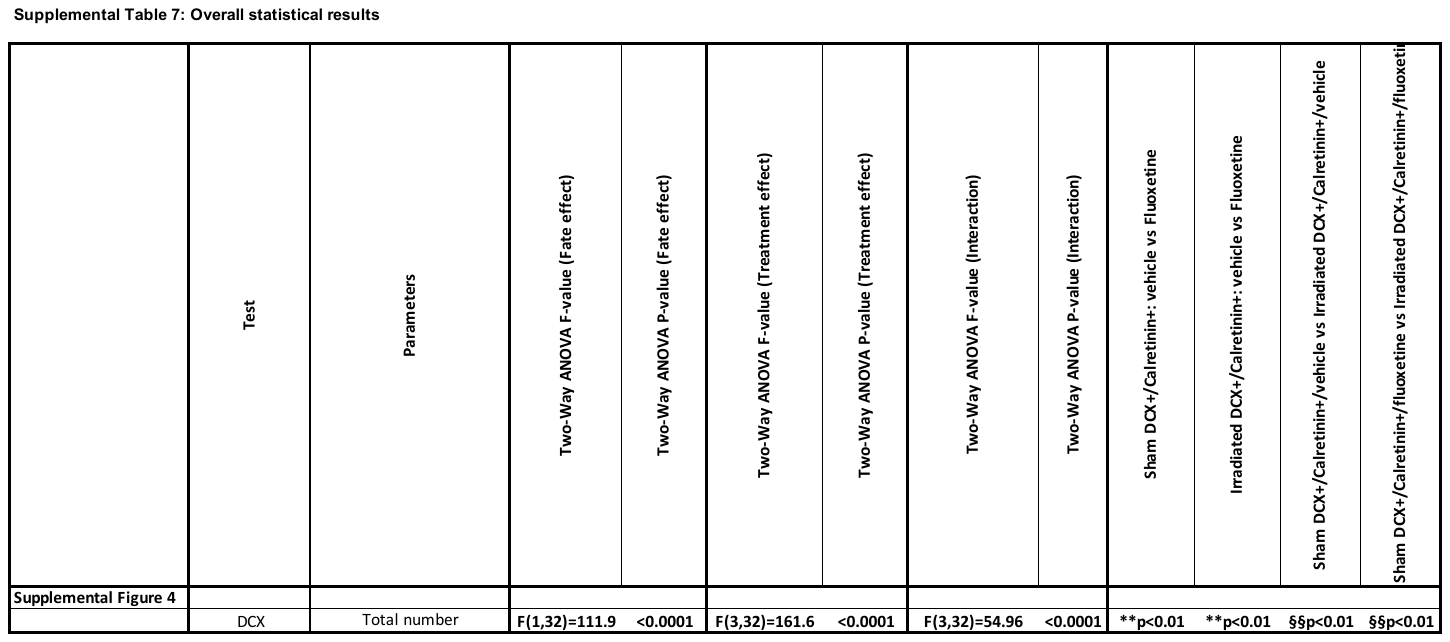

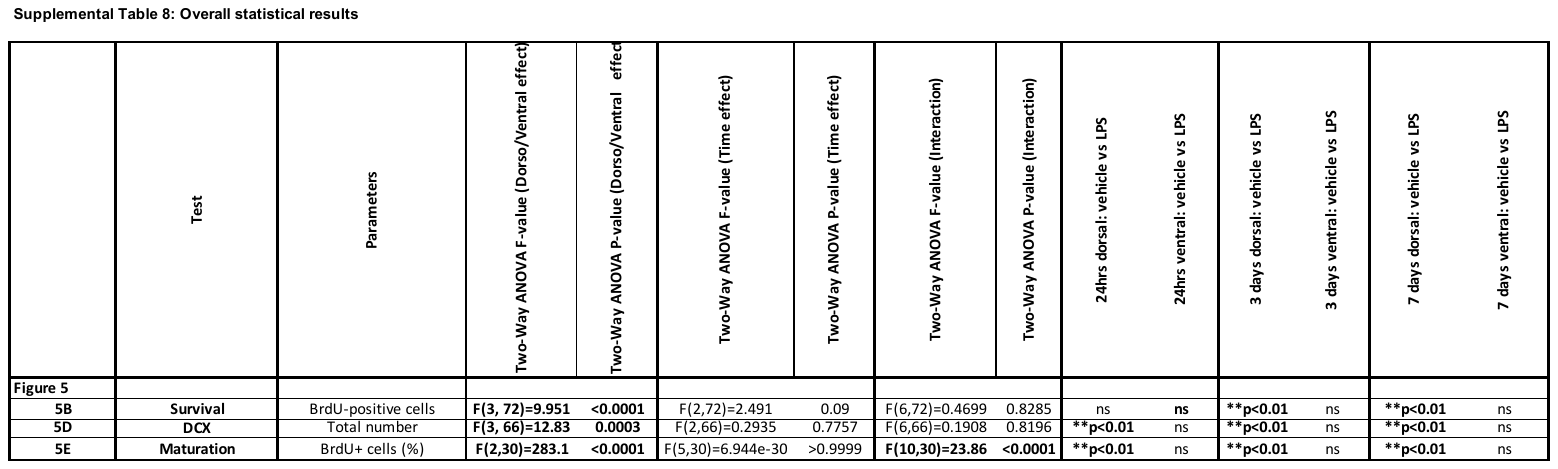

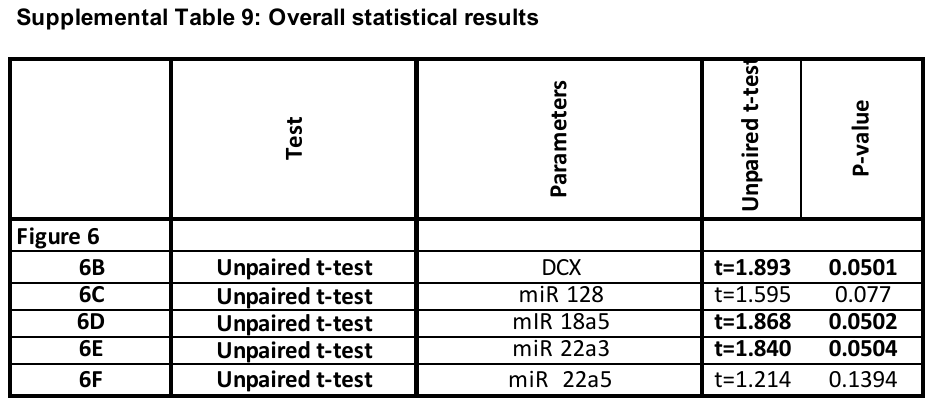

**SUPPLEMENTAL EXPERIMENTAL PROCEDURES**

**Animals**

**Study in β-arrestin 1 -/- KO mice**

β-arrestin 1 +/− were provided from Jean Martin Beaulieu (University of Toronto) (Saaber et al., 2019). Male heterozygous β-arrestin 1 +/− and heterozygous female mutant β-arrestin 1 +/− mice (age 4–6 months) were bred on a mixed S129/Sv × C57BL/6 genetic background at the University Paris-Saclay’s animal facility (Châtenay-Malabry, France). Resulting pups were genotyped by polymerase chain reaction (Saaber et al., 2019). Male heterozygous β-arrestin 1 +/+ KO and their littermates, 25–30 g body weight, were used to test the neurogenic effects of a chronic treatment with fluoxetine (18 mg/kg/day) (**Supplemental Figure 1**).

.

**REFERENCES**

Saaber, F., Schutz, D., Miess, E., Abe, P., Desikan, S., Ashok Kumar, P., Balk, S., Huang, K., Beaulieu, J.M., Schulz, S., and Stumm, R. (2019). ACKR3 Regulation of Neuronal Migration Requires ACKR3 Phosphorylation, but Not beta-Arrestin. *Cell Rep* 26**,** 1473-1488 e1479.
